# Transposon wave remodelled the epigenomic landscape in the rapid evolution of a novel X chromosome dosage compensation mechanism

**DOI:** 10.1101/2022.09.29.510218

**Authors:** David C.H. Metzger, Imogen Porter, Brendan Mobley, Benjamin A. Sandkam, Lydia J.M. Fong, Andrew P. Anderson, Judith E. Mank

## Abstract

Sex chromosome dosage compensation is a model to understand the coordinated regulation of transcription, however the advanced age of the sex chromosomes in model systems make it difficult to study how the complex regulatory mechanisms underlying chromosome-wide dosage compensation can evolve. The sex chromosomes of *Poecilia picta* have undergone recent and rapid divergence, resulting in widespread gene loss on the male Y, coupled with complete X chromosome dosage compensation, the first case reported in a fish. The *de novo* origin of a novel dosage compensation system presents a unique opportunity to discover new mechanisms of gene regulation and their evolutionary origins. By combining a new chromosome-level assembly of *P. picta* with whole-genome bisulfite sequencing and RNA-Seq data, we determine that the binding motif of Yin Yang 1 (YY1) is associated with male hypomethylated regions on the X, but not the autosomes. The male-specific hypomethylation of these motifs offers a putative model for male specific upregulation of genes on the X. These YY1 motifs are the result of a recent and rapid repetitive element expansion on the *P. picta* X chromosome, which is absent in closely related species that lack dosage compensation. Taken together, our results present compelling support that a disruptive wave of repetitive element insertions carrying YY1 motifs resulted in the remodeling of the X chromosome epigenomic landscape and the de novo origin of a new dosage compensation mechanism.

## Introduction

Some organisms with highly diverged sex chromosomes evolve complete dosage compensation to ameliorate the negative effects of haploinsuffiency.^1,2^ These complex mechanisms act across the entirety of the X chromosome and represent a key model to study the coordinated regulation of transcription^3^ through the integration of sex-specific genomic and epigenomic processes.^4^ Chromosome-wide dosage compensation mechanisms utilize sex-specific chromatin modifying complexes to balance expression between the haploid X and diploid autosomes in males, as well as to equalize expression of X-linked genes between the males and females^5,6^ regardless of their dosage sensitivity. This is in stark contrast to many other systems with highly diverged sex chromosomes where gene dose is only compensated for those loci that are dosage sensitive.^7^

Extensive analysis of dosage compensation model systems has provided extraordinary discoveries on the genomic and epigenomic mechanisms involved in the regulation of gene expression at the whole-chromosome level. In eutherian mammals, the long-non coding RNA from *Xist* is expressed early in development from the X inactivation centre and spreads along one X chromosome in female cells,^8,9^ ultimately leading to the transcriptional inactivation for most of the genes via the accumulation of epigenomic marks that are transmitted to all daughter cells.^10^ In *Drosophila*, dosage compensation of the X chromosome in somatic cells is achieved through the male-specific lethal (MSL) complex, which acetylates histone H4,^11^ resulting in hyper-expression of X-linked genes in males.^12^ Moreover, substantial insight on how dosage compensation mechanisms are established on neo-sex chromosomes has been gleaned from the spread of existing dosage compensation mechanisms to neo-sex chromosomes in *Drosophila*.^13^

However, our understanding of the evolutionary dynamics and mechanisms of sex chromosome divergence and the evolution of *de novo* dosage compensation mechanisms have thus far been limited due to the fact that the model systems for whole chromosome dosage compensation mechanisms are species with ancient sex chromosomes.^13^ Understanding the early stages of chromosome regulation in these models requires extensive extrapolation, and can make it difficult to differentiate cause from consequence.

Recently, the first case of complete sex chromosome dosage compensation in fishes was identified in *Poecilia picta*,^14^ and the close relative *P. parae.^15^ P. picta* and *P. parae* are closely related to the common guppy, *P. reticulata*, and the Endler’s guppy, *P. wingei*. These species all share the same sex chromosome yet they exhibit a remarkable diversity of sex chromosome divergence. Although several phylogenetic models have been suggested for the origin of the sex chromosomes and dosage compensation in these species,^16^ comparative phylogenetic and genomic methods^15,17,18^ suggest that the sex chromosomes diverged, and complete dosage compensation evolved in a short interval between 14.8-18.5mya.^15^ Importantly, the sex chromosomes of *P. reticulata* and *P. wingei* lack any evidence of dosage compensation,^14^ and karyogram data shows extensive homology between the X and Y relative to the highly degraded Y in *P. picta*.^19^ Taken together, this system creates a powerful comparative genomic framework to study the early evolution of sex chromosomes. Additionally, as the only known case of complete X chromosome dosage compensation in fish, it is unlikely that an existing mechanism of dosage compensation from a related species has been co-opted, presenting a unique opportunity to study the early *de novo* evolution of a novel dosage compensation mechanism.

In this study, we examine the genomic and epigenomic landscape of *P. picta* to identify sex-specific patterns consistent with X chromosome dosage compensation, embedded within a comparative framework with related species. To facilitate our analysis, we assembled and annotated a chromosome-level genome sequence of a single female using a combination of PacBio HiFi and HiC sequencing. Using whole genome bisulfite sequencing (WGBS), we identify sex-biased differentially methylated regions (DMRs) on the *P. picta* sex chromosome that are highly conserved among tissues. Using motif analysis of the DMR sequences, we identify enrichment of a putative DNA binding motif for the Yin Yang 1 (YY1) transcription factor. These YY1 motifs are highly enriched on the *P. picta* X chromosome as the result of a recent wave of repetitive element invasion, which is absent in closely related species that lack dosage compensation. We also find that these motifs are hypomethylated on the X in males but not in females, or on the autosomes of either sex. Interestingly, YY1 facilitates *Xist* inactivation of the X chromosome in eutherian mammals^20^ and the role of YY1 in *P. picta* appears to be a case of evolutionary convergence. Taken together, our results present compelling support for the remodeling of the X chromosome epigenomic landscape through a wave of repetitive element insertions carrying YY1 motifs. The male-specific hypomethylation of these motifs offers a putative model for male specific upregulation of genes on the X and the first piece of evidence towards elucidating the rapid *de novo* evolution of a dosage compensation mechanism in *P. picta*.

## Results

### *P. picta* genome assembly, annotation and statistics

Our assembly of the *P. picta* genome is 744 Mb in total length, with BUSCO completeness of 97.65% and an N50 of 33,053,486, representing a substantial improvement over previous genomic resources for the species.^21^ We recovered 23 chromosomes, numbered based on the syntenic comparison with the *P. reticulata* female genome assembly (Figure 1A),^22^ where chromosome 12 is the sex chromosome.^23,24^ It is worth noting that this numbering system differs from some previous reports.^19,25^ A total of 27,764 genes were annotated using RNA-seq data from head (with eyes removed), muscle, liver, testis, and ovary tissue, and 29.55% of the genome is classified as repeats (Supplemental Table 1).

**Figure 1.**
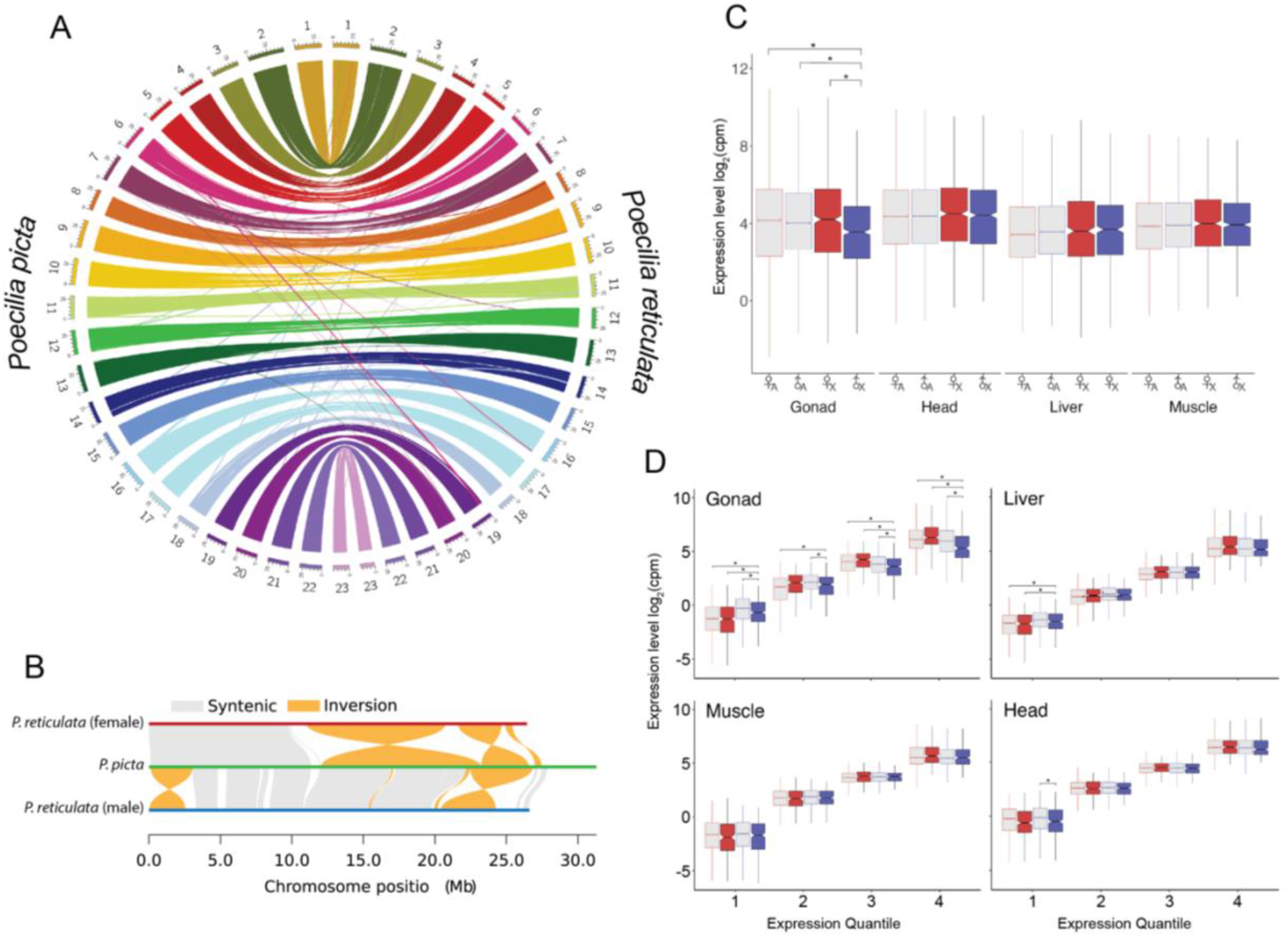
*Poecilia picta* genome assembly and analysis of tissue specific dosage compensation. A. Sequence synteny between the female *P. picta* and the female *P. reticulata* genomes. Chromosomes are labeled 1-23 and are differentiated by colour around the outer ring with *P. picta* chromosomes on the left and *P. reticulata* chromosomes on the right. The colour of the connecting lines indicates the chromosomal origin of the sequence from the *P. picta* genome. B. Chromosome 12 alignments between the female *P. reticulata*^22^ (red horizontal line), our *P. picta* (green horizontal line), and the male *P. reticulata*^26^ (blue horizontal line) genome assemblies. Grey segments connecting horizontal lines represent syntenic regions. Orange segments connecting horizonal lines indicate inverted regions. C. Comparison of autosomal (A, gray) and X chromosome (X, red or blue fill) gene expression values in female (red) and male (blue) gonad, head, liver, and muscle tissues. Gene expression levels presented as log2 counts per million (cpm) normalized to sequencing library size. The horizontal line of the box and whisker plot is the median, the box denotes the 25th and 75th percentile, and “whiskers” are 1.5 times the interquartile range. Significant difference (p<0.05) in gene expression between groups is indicated by an asterix (*), as determined by Bonferroni corrected Wilcoxon rank sum test. Complete list of p-values are available in Supplemental Table 3. D. Comparison of gene expression levels between autosomal and X chromosome genes in female and male gonad, head, liver, and muscle tissues in four expression quantiles. Colours and expression units are the same as in panel B. The complete list of p-values is available in Supplemental Table 4.

Synteny analysis between our *P. picta* and the short-read Illumina-based female *P. reticulata* genome assembly^22^ identified several large inversions and rearrangements (Supplemental Figure 1). In contrast, the *P. picta* genome is highly syntenic with the long-read male *P. reticulata*^26^ (Supplemental Figure 2), and *Xiphophorus helleri* (Supplemental Figure 3) genomes, suggesting that the apparent inversions in the female *P. reticulata* genome represent mis-assemblies. Of particular relevance for this study are the several large inversions on the X chromosome between *P. picta* and the male and female *P. reticulata* (Figure 1B). Linkage analysis of the sex determining region in *P. reticulata* has identified the sex determining locus to the distal end of chromosome 12,^23^ however there is little agreement in the orientation of this region between the three assemblies. This disparity could be caused by a lack of sequence complexity and an abundance of repetitive and duplicated sequences in this region, making it difficult to sequence and assemble consistently, particularly in the absence of long-read sequence data. The *P. picta* X chromosome (31.2Mb) also has ^~^4Mb at the distal end of the chromosome with very little homology to the X chromosome in *P. reticulata* (26.61Mb).

### Complete dosage compensation is observed in male somatic tissues but not testis

Following the split from *P. reticulata* and *P. wingei* ^~^18.4mya, the Y chromosome in the common ancestor of *P. picta* and *P. parae* rapidly diverged from the X over a period of 3-4my, leading to widespread gene loss and the rapid evolution of a complete dosage compensation mechanism.^14,15,25^ In contrast, the Y chromosome in *P. reticulata* and *P. wingei* has retained most of the coding content of the X and therefore there is little need for dosage compensation to evolve in these species.^14,27–29^

The effectiveness of dosage compensation mechanisms can vary between tissues.^30^ Previous studies of dosage compensation in *P. picta* and *P. parae* focused on the expression from samples consisting of mostly muscle tissue.^14,15^ To determine whether complete dosage compensation is equally effective across somatic and gonadal tissue, or whether dosage compensation is restricted to specific tissues, we first analyzed RNA-Seq data from male and female gonad, muscle, liver and head tissue. Across all somatic tissue, we observe that the average gene expression level from the X is not statistically different between males and females, and that the average gene expression level from the male X is not statistically different from male or female autosomes (Figure 1C; statistics available in Supplemental Table 2). However, the X expression was significantly lower in male gonad tissue compared to the average expression of male and female autosomal and female X genes in gonad. This is consistent with a ubiquitous complete dosage compensation mechanism acting in somatic tissues (Figure 1C, Supplemental Table 2).

The capacity of dosage compensation mechanisms to buffer the effects of gene dosage imbalances can be limited for highly expressed genes.^13,31^ To determine whether the efficiency of dosage compensation is affected by gene expression level, we analyzed dosage compensation patterns in four expression quantiles. Within each expression quantile, the average expression of sex chromosome genes in males is not significantly lower compared to male or female autosomal genes or female sex chromosome genes in somatic tissue, suggesting that dosage compensation is not affected by gene expression levels in *P. picta* (Figure 1D, Supplemental Table 4). Sex chromosome genes expressed in the head were significantly lower than autosomal genes for the lowest expression quantile in males, however the same pattern was also observed in females suggesting that this difference is not caused by the efficiency of dosage compensation in genes with low expression.

### The *P. picta* sex chromosome is differentially methylated in somatic tissues

Sex-specific epigenomic marks are commonly associated with facilitating sex-specific components of dosage compensation mechanisms, such as parent-of-origin specific differences in DNA methylation^32^ and maintenance of X inactivation in mammals,^33^ or histone acetylation by the MSL complex in *Drosophila*.^34^ We therefore examined male and female muscle, liver, and gonad tissues for sex-specific DNA methylation patterns using WGBS data. Whole genome DNA methylation values were similar in male and female muscle and liver tissue, while testis tissue was more highly methylated compared to ovary or somatic tissue (Supplemental Figure 4). Principal component analysis also revealed that DNA methylation patterns in testis are distinct from other tissues (Supplemental Figure 5). Analysis of DNA methylation values at the chromosome level revealed that CpG islands on the X chromosome are more highly methylated in both males and females compared to autosomal DNA methylation in all tissues (Supplemental Figure 6).

We next examined the WGBS data at the single nucleotide level to identify site specific patterns of sex-biased DNA methylation. Male and female gonads exhibit the greatest number of differentially methylated loci (DMLs) (744,852) and 79% of the DMLs are hypo-methylated in testis compared to ovary. DMLs between testis and ovary are evenly distributed throughout the genome (Supplemental Figure 7), with no enrichment on the X chromosome. We found relatively fewer sex-biased DNA methylation sites between male and female somatic tissues (9,575 DMLs in muscle; 7,536 DMLs in liver). In contrast to gonad tissue, sex-specific DNA methylation patterns in somatic tissue were highly enriched on the X chromosome (74% of DMLs in muscle and 82% of DMLs in liver compared to 5% in gonad).

The spatial distribution of CpG sites is not uniform throughout the genome. DMLs tend to occur in clusters forming regions of contiguous differential methylation profiles called Differentially Methylated Regions (DMRs). We therefore next analyzed the WGBS data for sex-biased DMRs. We found that the genomic distribution of DMRs is consistent with the distribution of DMLs (Figure 2, Supplemental Figure7–8). We found 1268 sex-biased DMRs between gonad tissues, 295 DMRs in muscle tissue, and 221 DMRs in liver tissue. The majority of DMRs in somatic tissues are hypomethylated in males (80% in muscle and 84% in liver) and are enriched on the sex chromosome (20% in muscle and 63% in liver), while the distribution of sex-specific DMRs in gonad were equally hypo- and hypermethylated in gonad tissue (53% hypomethylated and 47% hypermethylated in males), and were evenly distributed among autosomes and the sex chromosome (Figure 2, Supplemental Figure 7).

**Figure 2.**
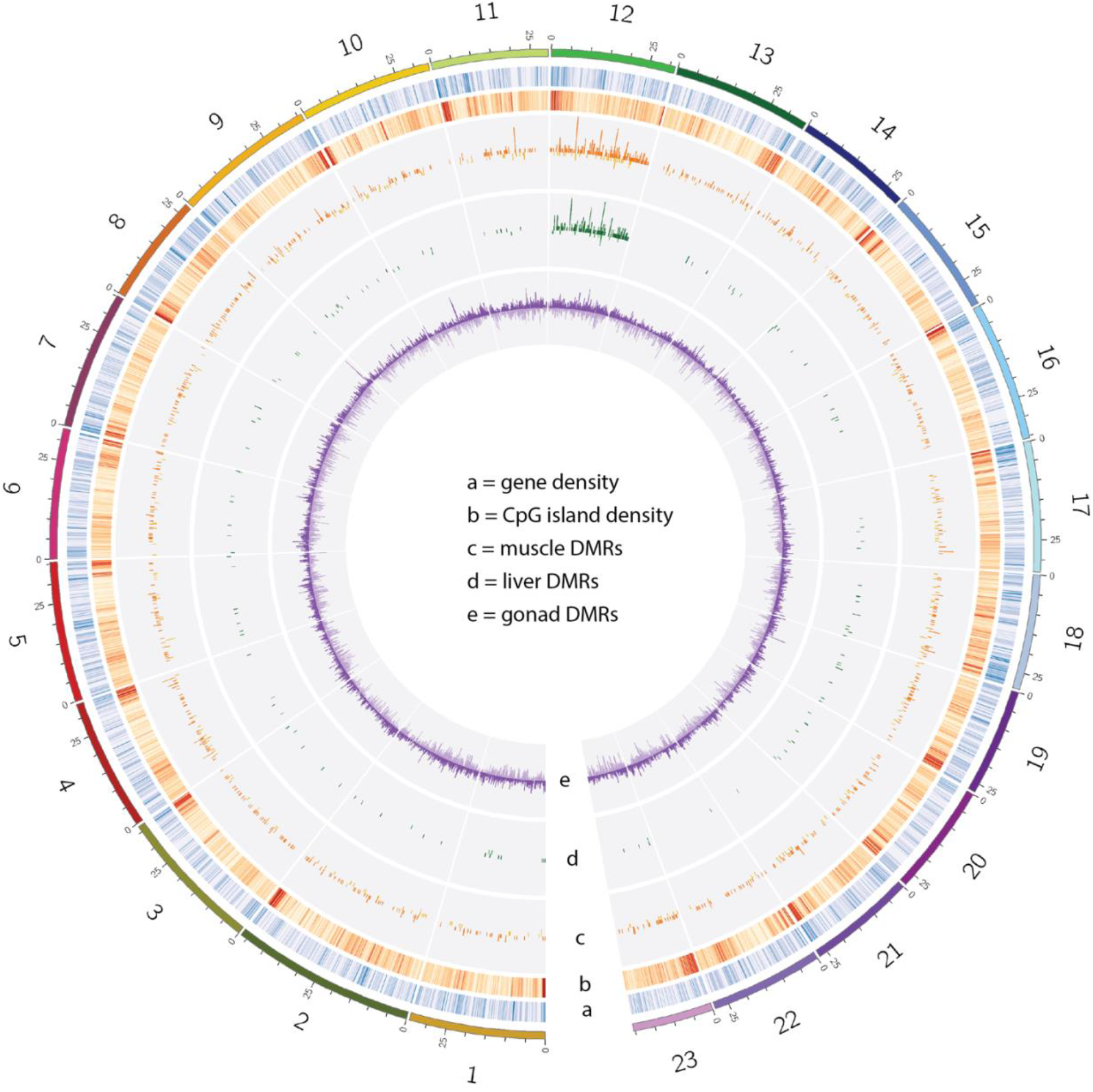
Sex-specific DNA methylation patterns in *P. picta*. Outer ring of circos plot indicates chromosome number and Mb position. Chromosome colours are the same as in Figure 1. Tracks a & b are heatmaps depicting gene (blue) and CpG island (red) densities, each in 100kb bins. Darker colors in the heatmap depict higher counts. Tracks c-d depict histograms of the number of differentially methylated regions (DMRs) in 100kb bins in muscle (orange), liver (green), and gonad (purple) tissues. Histogram bars extending outward represent DMRs that are hypomethylated in males while bars that extend towards the center of the plot represent DMRs that are hypomethylated in females. Chromosome 12 is the X chromosome.

Hypomethylated DMRs in promoter regions can activate the expression of nearby genes resulting in sex biased expression.^35^ Consistent with having a role in regulating gene expression, we found lower levels of DNA methylation in the promoter regions of highly expressed genes (Supplemental Figure 9). To determine whether DMRs on the sex chromosome are involve in gene-by-gene dosage compensation, we next examined DNA methylation levels of CpG islands located in the promoter regions of genes. We found no evidence for chromosome-wide sex-biased DNA methylation of gene promoters (Supplemental Figure 10), suggesting that the increased gene expression activity of male dosage compensation on the X chromosome is not regulated by DNA methylation on a gene-by-gene basis.

Some cases of sex chromosome dosage compensation are associated with heterochromatization.^36,38^ We therefore examined male and female *P. picta* karyograms for evidence of chromosome wide differences in heterochromatinization between male and female X chromosomes. Consistent with with karyograms of *P. picta* from a Trinidadian population (Nanda et al., 2022), we identified 46 acrocentric chromosomes, including one pair of heteromorphic XY sex chromosomes in males. We found the size of the X chromosome to be very similar between males and females (Supplemental Figure 11) suggesting that overall heterochromatization of the X chromosome does not differ substantially between the sexes.

### Genomic regions with sex-biased DNA methylation are enriched for YY1 motifs

We next examined the DNA sequences of the DMRs for the possibility of enriched regulatory elements. Analysis of DMR sequences on the sex chromosome identified 35 putative regulatory elements enriched in regions with sex-biased DNA methylation. The ten most significantly enriched motifs are presented in Table 1. The most significantly enriched regulatory element was the highly conserved DNA binding motif for the ubiquitously expressed YY1 transcription factor, which has a well-established role in sex-specific regulation of the *Xist* long non-coding RNA and dosage compensation in mammals.^20,37,39–41^

**Table 1.**
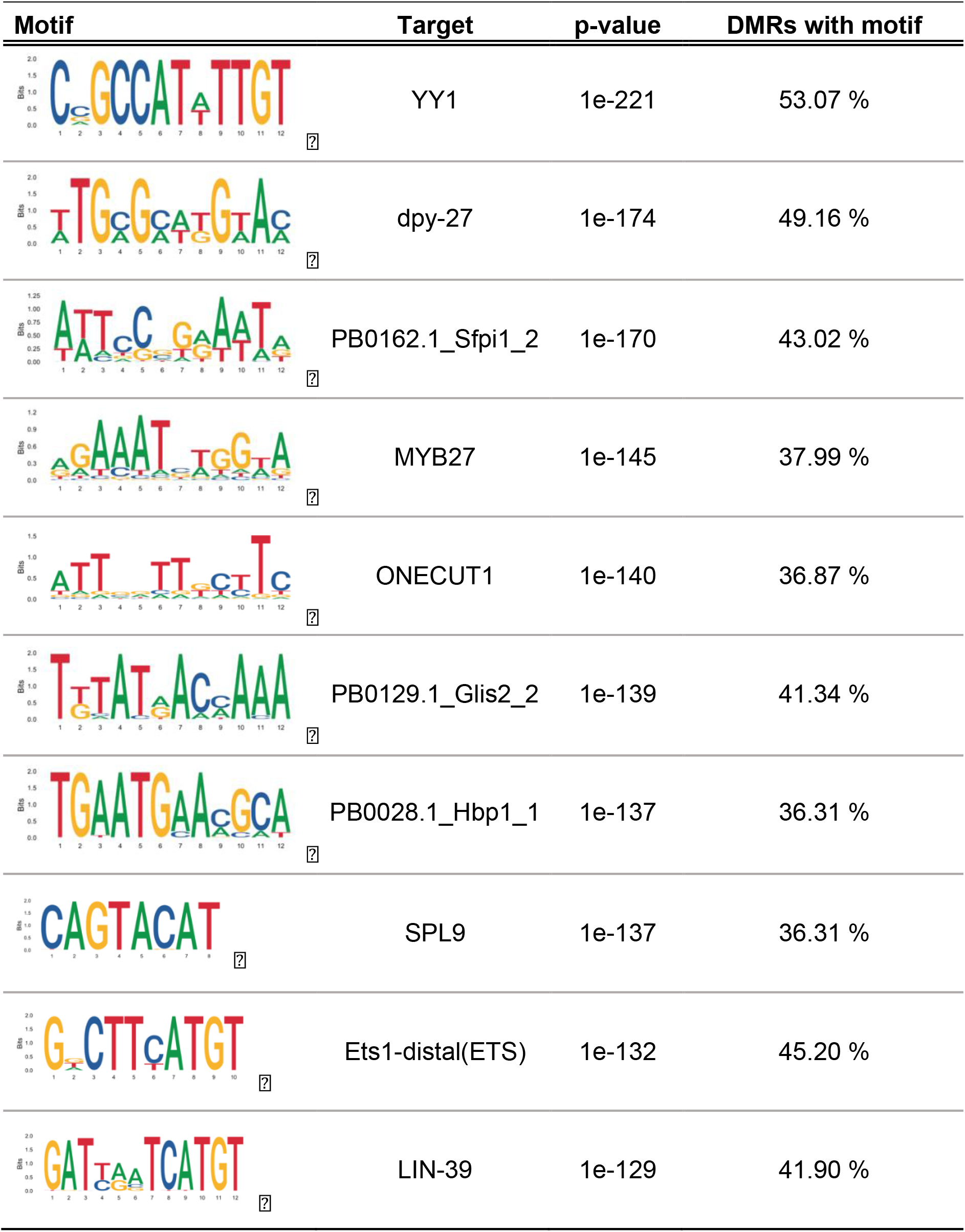
Motifs associated with DMRs on the *P. picta* X chromosome.

| Motif | Target | p-value | DMRs with motif |
| --- | --- | --- | --- |
|  | YY1 | 1e-221 | 53.07 % |
|  | dpy-27 | 1e-174 | 49.16 % |
|  | PB0162.1_Sfpi1_2 | 1e-170 | 43.02 % |
|  | MYB27 | 1e-145 | 37.99 % |
|  | ONECUT1 | 1e-140 | 36.87 % |
|  | PB0129.1_Glis2_2 | 1e-139 | 41.34 % |
|  | PB0028.1_Hbp1_1 | 1e-137 | 36.31 % |
|  | SPL9 | 1e-137 | 36.31 % |
|  | Ets1-distal(ETS) | 1e-132 | 45.20 % |
|  | LIN-39 | 1e-129 | 41.90 % |

Further analysis of the YY1 sequence motif revealed a dramatic abundance of YY1 motifs on the sex chromosome compared to the autosomes, likely resulting from the replication of three of the six possible motifs (CCGCCATATTGT, CGGCCATTTTGT, and CAGCCATTTTGT) on the sex chromosome relative to the autosomes (Figure 3A). These motifs are highly conserved compared to mammalian YY1 motifs, and one in particular (CCGCCATATTGT) is identical to the YY1 binding motif of the *Xist* loci (CCGCCATnTT) which is regulated by sex-biased DNA methylation and is involved in mammalian dosage compensation.^42^

**Figure 3.**
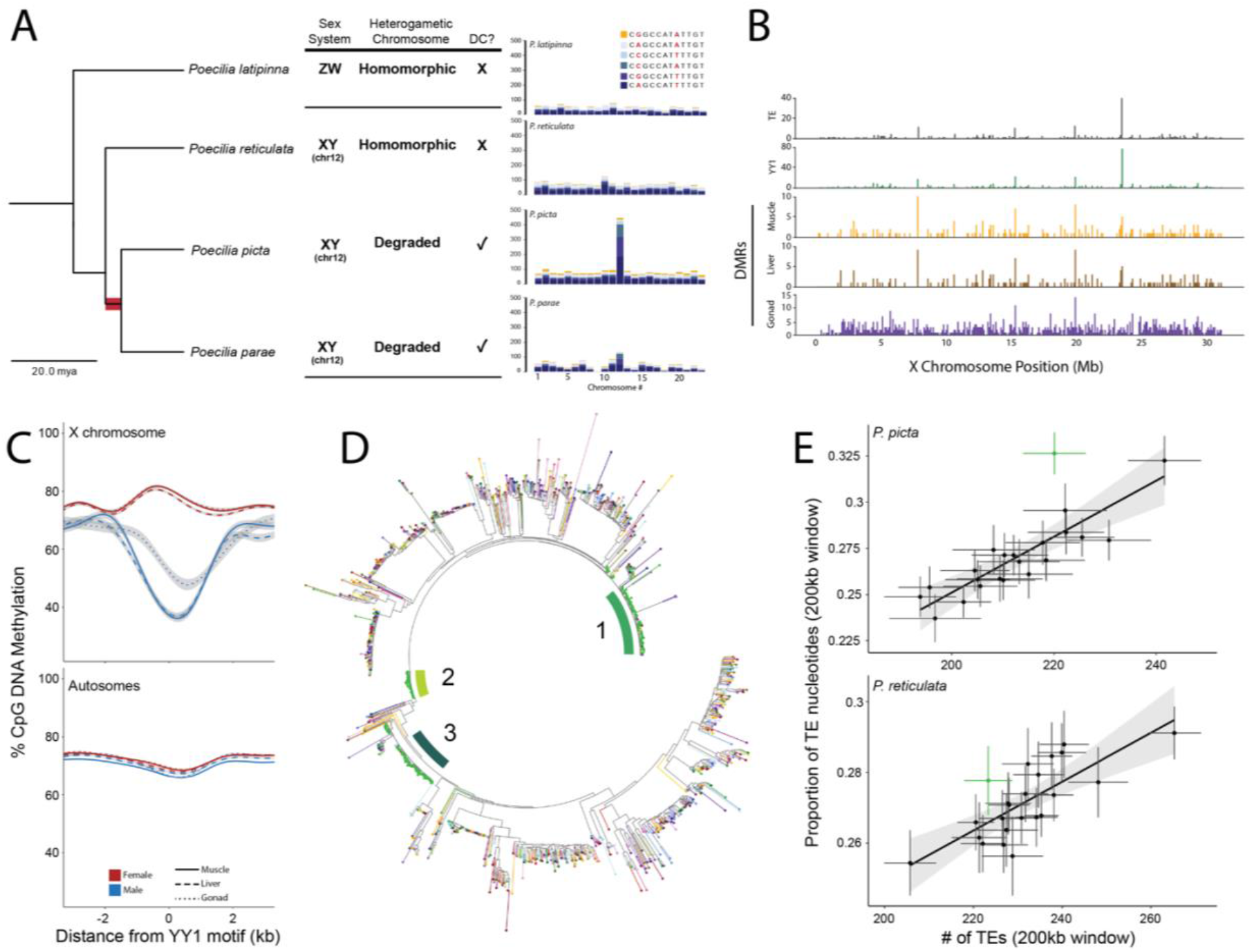
YY1 motif accumulated on the *P. picta* X chromosome from a wave of TE replication and exhibits sex-specific DNA methylation. A. Phylogeny of *Poecila* species depicting the diversity of XY heteromorphism between *P. reticulata, P. picta*, and *M. parae* and the ^~^3.7-Myr interval (shaded in red) for when the Y chromosome degenerated and dosage compensation arose in the common ancestor of *P. picta* and *P. parae*. Barplots depict the chromosomal distribution of YY1 motifs. Colours represent the different YY1 motif sequences. Nucleotides highlighted in red represent the nucleotide positions that vary between motifs. The phylogeny and divergence times are taken from The Fish Tree of Life,^97^ and adapted from^25^ and^15^. B. Bar plots depicting the number of YY1 motif containing TEs (gray), YY1 motifs (green), male specific hypomethylated regions in muscle (orange), liver (brown), and gonad (purple) tissue in 100kb bins along the X chromosome. C. DNA methylation levels of genomic regions containing YY1 motifs +/- 2500bps in females (red) and males (blue) on the X (upper panel) and autosomes (bottom panel) presented as a loess regression with 95% confidence intervals in gray. Tissue specific DNA methylation indicated by line type; muscle = solid line, liver = dashed line, gonad = dotted line. Zero marks the location of a YY1 motif. D. Circular phylogeny of repetitive sequences containing at least one YY1 motif. Chromosomes are indicated by different colours as in Figures 1 and 2 with the X chromosome in green. Green bars indicate the three clades where sequences are almost exclusively derived from the X chromosome. E. Relationship between the length and number of repetitive elements on each chromosome. Each point represents mean values in 200kb bins for each chromosome (X = green). Length is calculated as the total length of repetitive elements in a 200kb bin. Length represents the sum of the lengths for every repetitive element in a 200kb bin. Confidence intervals are standard error. Count represents the total number of repetitive elements in 1Mb bins. Shaded region is the 95% confidence interval line around the linear regression (solid black line).

To determine whether the enrichment of YY1 motifs on the X chromosome is present in other Poeciliids, we analyzed genomic sequences from several related species (Figure 3A). YY1 sequence motifs were enriched on the X chromosome of the sister species *P. parae*, which also exhibits complete X chromosome dosage compensation.^15^ We found no enrichment of YY1 motifs on the X chromosome of *P. reticulata*, which shares the same X chromosome but lacks substantial Y degeneration or dosage compensation^14^ or *P. latippina*, which has a ZW sex determining mechanism located on a different chromosome from *P. picta* ^14,43^ (Figure 3A).

Comparison of the YY1 motif distribution along the X chromosome revealed a striking similarity to the distribution of DMRs, particularly those in somatic (i.e. dosage compensated) tissues (Figure 3B). YY1 motifs have been shown to be regulated by differential DNA methylation patterns in other species^37,44^ and is known to play a role in *Xist* mediated X chromosome inactivation in eutherian mammals.^20,40,45^ To determine whether sex-biased DNA methylation patterns in *P. picta* could result in sex-biased YY1 binding to the regulatory motif, we analyzed DNA methylation levels around the YY1 motif on the sex chromosome and on the autosomes separately. We find that YY1 motifs on the X chromosome are hypomethylated in all male tissues relative to females (Figure 3C). The percent difference in DNA methylation of YY1 motifs on the sex chromosome were similar in muscle (50%) and liver (48%) tissue. DNA methylation of YY1 motifs in testis were also hypomethylated but to a lesser extent (29%) compared to somatic tissues. In contrast, we observe no difference between male and female DNA methylation levels for YY1 motifs on the autosomes (Figure 3C).

To determine whether male-specific hypomethylation of YY1 motifs results in a localized hyper-expression of genes, we also analyzed gene expression levels in the immediate vicinity of YY1 motifs on the X chromosome and on autosomes. We found no difference in the expression of genes around YY1 motifs between males and females in any tissue (Supplemental Figure 12). These data demonstrate that methylation of YY1 sites does not regulate expression on a gene-by-gene basis and suggests a putative dosage compensation mechanism mediated by YY1 binding affinity and long-range control of gene expression. Annotation of the *P. picta* genome identified two YY1 orthologs, ANN22864 on ch21 and ANN18583 on chr22. Both YY1 orthologs are ubiquitously expressed in head, muscle, liver, and gonad tissues (Supplemental Figure 13) and their DNA binding domain, consisting of four C2H2-type zinc fingers, is identical to the mammalian YY1 DNA binding domain and the DNA binding domain of other fishes (Supplemental Figure 14).

### Recent expansion of repetitive elements containing YY1 motifs

YY1 motifs can regulate the expression of LINE-1 transposable elements (TE),^46,47^ providing a potential means for their replication and expansion throughout the genome. To test whether the enrichment of YY1 motifs on the *P. picta* X chromosome can be attributed to TE replication, we isolated repetitive regions of the genome, and identified those containing at least one YY1 motif for phylogenetic analysis (Figure 3D). This phylogenetic analysis revealed that most of these unannotated repetitive sequences cluster with sequences from across the autosomes, but three distinct clusters contain repetitive sequences that are present, almost exclusively, on the X chromosome. The short internal branch lengths of these three clusters and their strict localization on the X strongly suggest that they originated from recent TE expansions. To further examine the repetitive element profile of the *P. picta* genome, we calculated Kimura distance values using 200kb sliding windows for each chromosome. Of the 23 chromosomes, the sex chromosome contains an anomalous abundance of longer repeats (Figure 3E) with low Kimura distance values (Supplemental Figure 15) consistent with a recent expansion of repetitive elements on this chromosome, which may partially account for the size discrepancy between the X chromosome assembly size between *P. picta* (31.3Mb) and *P. reticulata* (26.6Mb) (Figure 1B). Taken together, these results suggest that a recent expansion of transposable elements along the X chromosome in *P. picta* produced the rapid accumulation of the YY1 transcription factor motifs.

## Discussion

Understanding how dosage compensation mechanisms originate and evolve is important for understanding the molecular and evolutionary processes that regulate chromosome structure and gene regulatory processes.^3,48^ However, it remains unclear how these complex adaptations to Y chromosome degeneration can arise, as the sex chromosomes of model systems are ancient, making it difficult to extrapolate the initial stages of the *de novo* origin of dosage compensation, and obscuring cause from consequence. From our high quality, chromosome-level assembly of *P. picta*, we were able to identify key elements of the genomic and epigenomic architecture that led to the recent and rapid evolution of a novel dosage compensation mechanism. Our results support the model that a recent wave of transposon replication led to a sex-chromosome specific accumulation of YY1 DNA binding motifs and that sex-biased DNA methylation promotes male specific activation of the YY1 motifs as a key component in establishing dosage compensation in *P. picta* (Figure 4).

**Figure 4.**
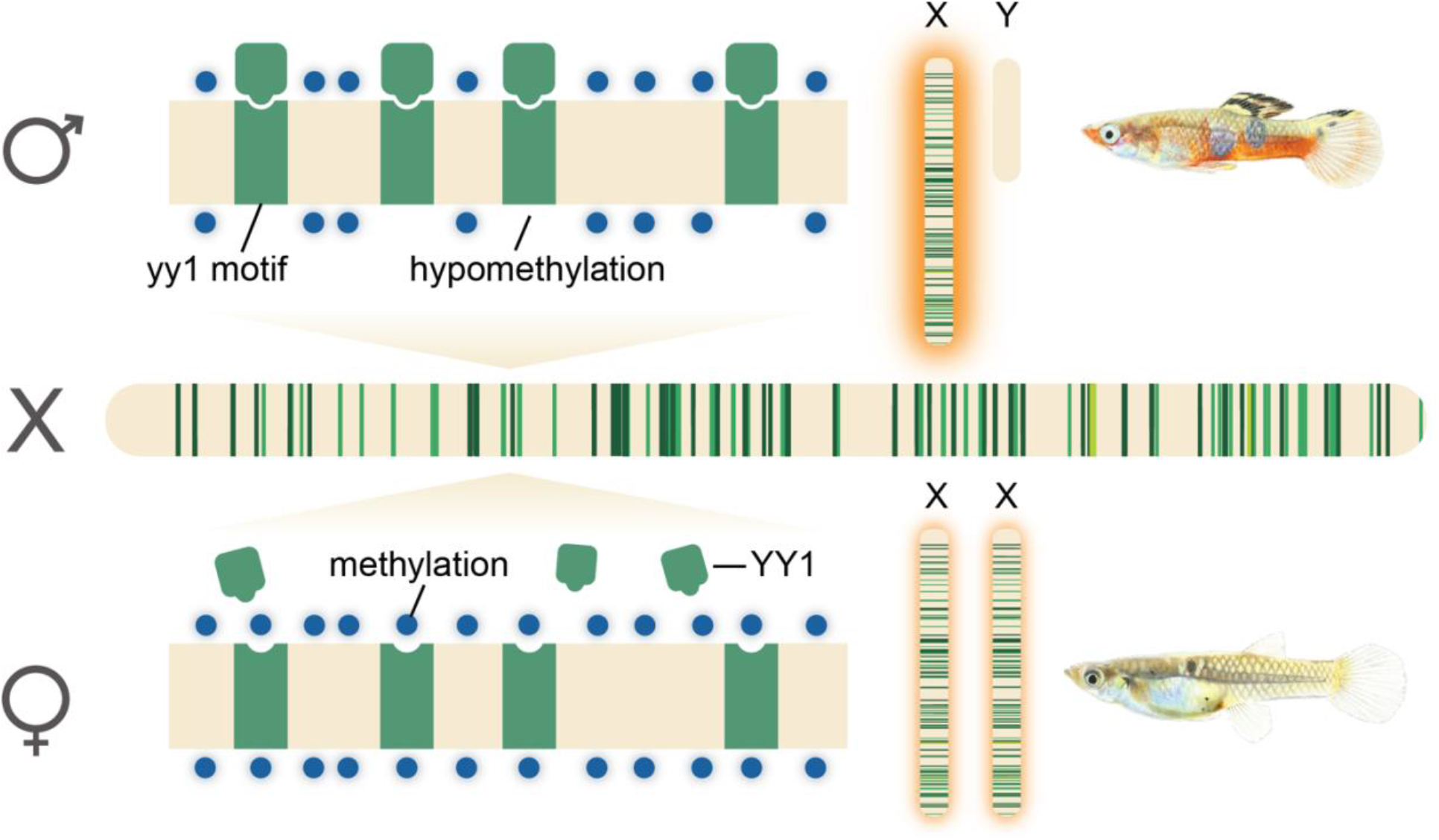
Proposed model for complete X chromosome compensation in *P. picta*. Hypomethylation of YY1 DNA binding motifs in males promotes male-specific binding of the YY1 transcription factor and the global upregulation of genes on the X chromosome in males. Hypermethylation of YY1 DNA motifs in females prevents YY1 binding and the hyperexpression of X chromosome genes in females. Green bars indicate position of YY1 motifs along the X chromosome and correspond to the clade of repetitive elements that the YY1 motif is located in based on colors in Figure 3.

### YY1 and dosage compensation in Poecilia picta

YY1 is a ubiquitously expressed^49,50^ DNA-binding zinc-finger transcription factor ^51,52^ that has been shown to selectively bind to hypo-methylated DNA sequences.^53,54^ In mammals, YY1 shares several characteristics with the conserved CCCTC-binding factor protein (CTCF). Both CTCF and YY1 are ubiquitously expressed transcription factors that function as activators, repressors, chromatin insulators, and play important roles in imprinting.^41^ CTCF protein has a well-established role in gene regulation and the formation of 3D chromatin organization such as topologically associated domains in mammals.^55^ These gene expression domains are often demarcated by the formation of newly derived CTCF motifs resulting from a recent expansion of repetitive elements.^56^ Analogous to CTCF-CTCF interactions, YY1 dimerization has also been shown to promote the formation of DNA loops, forming insulated regions of and facilitating enhancer-promoter interactions.^39,49,57,58^

Our results are consistent with a recent expansion of YY1 motifs and the compartmentalization of the X chromosome in the common ancestor of *P. picta* and *P. parae*. These data suggest a convergent model for the evolution of regulatory domains and novel dosage compensation mechanisms through a transposable element mediated expansion of transcription factor motifs.^59,60^ Additionally, allele specific expression patterns in *P. picta* and *P. parae* strongly suggest that dosage compensation in these species is achieved through a male-specific hyperexpression of the X chromosome,^14,15^ a process comparable to dosage compensation in *Drosophila*.^34^ Our study suggests that male-specific hyperexpression of the X chromosome is achieved by male-specific hypomethylation of YY1 motifs on the X chromosome, thereby facilitating male-specific activation of these motifs and modifying expression of the entirety of the X chromosome. Importantly, transcription of YY1 regulatory elements produces a positive-feedback loop that contributes to the stability of gene expression programs ^61^ which could provide a mechanism to maintain sex-specific activity of YY1 motifs once they are established.

Sex-specific regulation of YY1 elements is well established, and plays a pivotal role in genomic imprinting and sex-specific regulation of the *Xist* long non-coding RNA involved X chromosome silencing in mammals.^20,37,39–41^ In the context of mammalian dosage compensation, YY1 binding is differentially regulated by the sex-specific epigenetic landscape between the active and inactive X in females. Our data supports the convergent evolution of a YY1 mediated dosage compensation mechanism regulated by the epigenomic landscape in *P. picta*. While the exact role of YY1 in *P. picta* is still unclear, future studies will provide key insights into the role of YY1 and the regulation of chromosome structure, enhancer-promoter interactions, and as a nucleation factor for the binding and spreading of dosage compensation machinery^20,39,41^ on the *P. picta* X chromosome. Although some cases of sex chromosome dosage compensation are associated with heterochromatization,^36,38^ morphological analysis of metaphase spreads from males and females in *P. picta* showed no clear difference between male and female X chromosome morphology. Comparison of male and female X chromosomes using higher resolution methods, such as C-banding which can selectively stain heterochromatin-rich regions of individual chromosomes, may be more effective at identifying sex-specific differences in X chromosome chromatin organization^62,63^, or it may be that X chromosome dosage compensation is achieved without differential heterochromatinization between males and females, as is the case in *Drosophila*.^67^

### TE replication and sex chromosome evolution

Transposable elements are major innovators of genome evolution and speciation. Replication and expansion of genomic regions via transposable element activity is an important process that contributes to the generation of genomic diversity such as the replication and formation of new TF binding sites.^56,64,65^ In some cases, notably the case of the *Drosophila* neo sex chromosomes,^66^ existing mechanisms of dosage compensation can spread with the expansion of binding motifs.^67^ However, many of the genes involved in the formation and spreading of the *Drosophila* MSL complex are not present in fish genomes.

Younger, potentially active TEs are silenced by epigenetic processes such as DNA methylation.^68,69^ In particular, LINE-1 retrotransposition is regulated by DNA methylation of YY1 transcription factor motifs.^46,47,70^ Furthermore, the copy and paste mechanism through which the expansion of retrotransposons occurs means that once a YY1 motif is established in a TE, it has the potential to spread throughout the genome. This provides a potential positive feedback loop for the expansion of TEs bearing YY1-motifs in hypomethylated regions.

We have identified a recent repeat-associated expansion YY1-binding motifs on the *P. picta* X chromosome, suggesting a potential model for the rapid evolution of dosage compensation. The X-chromosome enrichment of YY1 motifs and YY1-bearing TEs is absent from the syntenic chromosomes of *P. reticulata* and *P. latippina*, which both lack dosage compensation. This phylogenetic distribution suggests that the expansion occurred in the common ancestor to *P. picta* and *P. parae* ^~^18.4mya in tandem with the rapid evolution of dosage compensation over 3-4myr^15^ (Figure 3A). Various models have been proposed for the evolutionary history of sex chromosomes in the clade of Poeciliids encompassing *P. reticulata, P. wingei, P. picta*, and *P. parae*.^14,16,18^ However, all available evidence suggests that the sex chromosome arose on in the ancestor to *P. reticulata, P. wingei, P. picta* and *P. parae*, roughly 20 mya.^15,17,18,25^ Based on this model, extensive Y chromosome degradation and dosage compensation evolved in the common ancestor of *P. picta* and *P. parae* after the split from *P. reticulata* and *P. wingei*.

Our data here are also consistent with the recent origin of the sex chromosomes, roughly 20mya. The enrichment of YY1 motifs is specific to the X chromosome in *P. picta* and *P. parae*, the two species with complete sex chromosome dosage compensation. Because Y chromosome degeneration is facilitated by the evolution of dosage compensation^71^ it is reasonable that the extensive sex chromosome divergence we observe in *P. picta* and *P. parae* was associated with the YY1-associated dosage compensation mechanism. If the divergence and dosage compensation mechanism were much older and simply associated with sex chromosome turnover events in other species, as has been suggested by some,^16^ we would expect to observe the enrichment of YY1-motif bearing TEs on the syntenic regions in other related species, which we do not. Moreover, phylogenetic analysis of repetitive element sequences containing YY1 motifs supports the model of a recent transposable element wave that resulted in the accumulation of YY1 elements on the X, again consistent with a recent origin of dosage compensation.

## Conclusion

Understanding the structure, function, and origins of genomic processes is fundamental to understanding the mechanisms of genomic evolution. *Poecilia* is the only known clade of fishes with complete X chromosome dosage compensation. Aside from mammals, the only other vertebrate for which a complete dosage compensation mechanism has been proposed is in *Anolis* lizards,^72^ however this mechanism remains unknown. The extensive phylogenetic distance to other known mechanisms of whole chromosome dosage compensation make co-option of these mechanisms in *P. picta* highly unlikely. The dosage compensation system in *P. picta* therefore likely evolved largely *de novo* from a suite of genes and regulatory elements that acquired a novel role to selectively target and modulate expression of the X chromosome in a sex-specific manner. Here we present evidence for an elegant strategy to rapidly evolve a novel dosage compensation system through the recruitment of DNA binding motifs of a ubiquitously expressed transcription factor via waves of transposable element expansion coupled with sex-specific regulation of the YY1 regulatory motif via DNA methylation. These data support the convergent evolution of YY1 as key component of dosage compensation between mammals and Poeciliids. However, the putative dosage compensation mechanisms described here represents a novel function of YY1, as fishes lack many of the genes and regulatory systems involved in mammalian and *Drosophila* dosage compensation mechanisms.

## Materials and Methods

### Fish Collection and Sampling for Genome Sequencing

Animals used in this study are from a laboratory-reared population of *P. picta* collected initially in the Spring 2019 from natural populations in Suriname and brought to the University of British Columbia (Vancouver, BC, Canada) aquatics facility, where they were kept in 20 L glass aquaria on a 12:12 day:night cycle at 26 °C and 13ppt salinity (Instant Ocean Sea Salt), and fed Hikari Fancy Guppy pellets supplemented with live brine shrimp daily. Prior to tissue sampling, individuals were euthanized using a lethal overdose of MS-222. For genome sequencing, muscular tail tissue was taken from the anal pore to the base of the pectoral fin from a single adult female. Tissue was immediately flash frozen in liquid nitrogen and stored at −80 °C.

### PacBio Library and Sequencing

Frozen tissue samples were shipped overnight on dry ice to the Dovetail Genomics Production Lab (Scotts Valley, CA) for Dovetail Omni-C Library Preparation and Sequencing. DNA samples were quantified using Qubit 2.0 Fluorometer (Life Technologies, Carlsbad, CA, USA). The PacBio SMRTbell library (^~^20kb) for PacBio Sequel was constructed using SMRTbell Express Template Prep Kit 2.0 (PacBio, Menlo Park, CA, USA) using the manufacturer recommended protocol. The library was bound to polymerase using the Sequel II Binding Kit 2.0 (PacBio) and loaded onto PacBio Sequel II. Sequencing was performed on PacBio Sequel II 8M SMRT cells.

### De novo sequence assembly

The wtdbg2 long-read assembler^73^ was run with the following parameters: genome_size 0.7g --read_type sq --min_read_len 20000 --min_aln_len 8192. Blobtools v1.1.1^74^ was used to identify potential contamination in the assembly based on blast (v2.9) results of the assembly against the NT database. A fraction of the scaffolds were identified as contaminant and were removed from the assembly. The filtered assembly was then used as an input to purge_dups v1.1.2^75^ and potential haplotypic duplications were removed from the assembly.

### Dovetail Omni-C library preparation and sequencing

To generate the Omni-C libraries, chromatin was fixed with formaldehyde in the nucleus and then extracted. The extracted chromatin was then digested with DNAse I. Chromatin ends were repaired and ligated to a biotinylated bridge adapter followed by proximity ligation of adapter containing ends. Crosslinks were then reversed and the DNA purified. Purified DNA was treated to remove biotin that was not internal to ligated fragments. Sequencing libraries were generated using NEBNext Ultra enzymes and Illumina-compatible adapters. Biotin-containing fragments were isolated using streptavidin beads before PCR enrichment of each library. The library was sequenced on an Illumina HiSeqX platform to produce approximately 40x sequence coverage. Reads with MQ>50 were then used for scaffolding.

### Scaffolding the Assembly with HiRise

The input de novo assembly and Dovetail OmniC library reads were used as input data for HiRise, a software pipeline designed specifically for using proximity ligation data to scaffold genome assemblies.^76^ Dovetail OmniC library sequences were aligned to the draft input assembly using bwa.^77^ The separations of Dovetail OmniC read pairs mapped within draft scaffolds were analyzed with HiRise to produce a likelihood model for genomic distance between read pairs. This model was used to join scaffolds, to identify and break putative misjoins, and to score prospective joins. Genome completeness was estimated based on Benchmarking Universal Single-Copy Orthologs (BUSCOs) against the eukaryota_odb10 database.^78^

### Genome annotation

Coding sequences from *P. formosa, P. latipinna, P. mexicana, P. reticulata* and *X. maculatus* were used to train the initial ab initio model for *P. picta* using the AUGUSTUS software (version 2.5.5). Six rounds of prediction optimisation were done with the software package provided by AUGUSTUS. The same coding sequences were also used to train a separate ab initio model for *P. picta* using SNAP (version 2006-07-28). RNAseq reads were mapped onto the genome using the STAR aligner software (version 2.7) and intron hints generated with the bam2hints tools within the AUGUSTUS software. MAKER, SNAP and AUGUSTUS (with intron-exon boundary hints provided from RNA-Seq) were then used to predict for genes in the repeat-masked reference genome. To help guide the prediction process, Swiss-Prot peptide sequences from the UniProt database were downloaded and used in conjunction with the protein sequences from *P. formosa, P. latipinna, P. mexicana, P. reticulata* and *X. maculatus* to generate peptide evidence in the Maker pipeline. Only genes that were predicted by both SNAP and AUGUSTUS softwares were retained in the final gene sets. To help assess the quality of the gene prediction, AED scores were generated for each of the predicted genes as part of the MAKER pipeline. Genes were further characterised for their putative function by performing a BLAST search of the peptide sequences against the UniProt database. tRNA were predicted using the software tRNAscan-SE (version 2.05).

### Sequencing the mitochondrial genome

To sequence the mitochondrial genome, we extracted liver and brain from a male individual and enriched for mitochondria using a Qproteome Mitochondria Isolation Kit (Qiagen). DNA was extracted using a QIAamp DNA Micro kit (Qiagen) and sequenced on a MiSeq as 250bp PE reads. Reads were trimmed using Trimmomatic^79^ resulting in 1.2 million sequences, which were de novo assembled using Geneious Prime^®^ (v.2022.2.1 Biomatters Ltd.). The resulting mitochondrial sequence was aligned sequences of close relatives (*P. parae, P. reticulata* and *P. formosa*) and annotations were lifted over. A 245bp insertion in the D-Loop of *P. picta* was confirmed using 3 sanger reactions to sequence the 1.7kb region spanning the insertion.

### Synteny analysis

Sequence synteny was compared between the *P. picta* genome and the genomes of *P. reticulata* (female Genbank accession# GCA_000633615.2 and male Genbank accession # GCA_904066995.1) and *X. hellerii* (Genbank accession# GCA_003331165.2). Syntenic regions were visualized using the output from satsuma2 in Circos.^80^ Synteny between the *P. picta* and *P. reticulata* X chromosome was visualized using plotsr.^81^ YY1 amino acid sequences were obtained from Ensembl (human=ENSP00000262238, mouse=ENSMUSP00000021692, pig=ENSSSCP00000053317, stickleback=ENSGACP00000011347, medaka=ENSORLP00000019577). Alignments between mammalian and fish YY1 amino acid sequencers was performed in Geneious (v.2022.2.1 Biomatters Ltd.).

### Mitotic chromosome preparation

Cytogenetic analysis was conducted on adult fish (two male, two female) from the *P. picta* population described above. Mitotic chromosome preparations were obtained from gill arches according to.^82^ Briefly, live specimens were exposed to a 0.01% colchicine solution for seven hours before being euthanized with a lethal overdose of MS-222 and dissected. Extracted organs were incubated in 0.4% KCl solution for 25-45 minutes and fixed with three changes of freshly prepared 3:1 ethanol : acetic acid fixative. Organs were minced in 50% acetic acid and mounted onto slides warmed to approximately 45°C, then stained for 10 minutes in a 5% Giemsa solution with phosphate buffer pH 6.8 and air-dried. Slides were visualized on a Nikon Eclipse Ti-S inverted microscope (100x/1.30NA oil objective) equipped with a Veroptics IMX174 color camera (or cooled monochrome equivalent) with Micro-Manager software (MMStudio version 2.0.0, MMCore version 10.1.1). For each individual specimen, all visible chromosome spreads were photographed and evaluated for completeness. A minimum of eight high-quality images per specimen were selected for further analysis. Captured images were edited and analyzed with Adobe Photoshop (version 23.5.1).

### Karyotype analysis

Chromosomes were sorted by length, grouped by morphology into homologous pairs, and classified according to.^83^ All chromosomes prepared from female specimens were successfully paired while male spreads consistently contained two chromosomes that could not be paired in this manner. The two heteromorphic chromosomes in the male karyotype were designated as sex chromosomes based on morphological and size similarity of the larger unpaired chromosome to the female X. For each chromosome spread, a relative length index was calculated for the X chromosome(s) by dividing the length of the X chromosome(s) by the mean length of the chromosomes in the longest pair (i.e., pair 1).

### RNA-seq sampling and analysis

Sampling of adult fish was performed as described above. Liver, head, and gonad tissue was sampled for RNA-seq analysis and genome annotation. A total of nine males and nine virgin females were sampled. For each tissue we pooled three individuals of the same sex to create a total of three non-overlapping male and three female sample pools for library preparation and sequencing. Frozen tissue samples were shipped on dry ice to Genewiz (South Plainfield, NJ) for RNA purification, mRNA library preparation and sequencing.

Total RNA extraction was done using the QIAGEN RNeasy Plus Kit following manufacturer protocols. Total RNA was quantified using Qubit RNA Assay and TapeStation 4200. Prior to library prep, RNA samples were DNase treated followed by AMPure bead clean up and QIAGEN FastSelect HMR rRNA depletion. Library preparation was done with the NEBNext Ultra II RNA Library Prep Kit following manufacturer protocols. Libraries were then sequenced on the NovaSeq6000 platform in 2 × 150 bp configuration. RNA-seq data from muscle tissue (SRA accession #’s SAMN29631966, SAMN29631977, SAMN29631978, SAMN29631989, SAMN31093659)^18^ was also used for genome annotation and RNA-seq analysis.

Sequencing data was quality filtered and Illumina adapters were trimmed using Trimmomatic v0.36.^79^ Reads were scanned with a 4-base sliding window and discarded when the average phred score was <15. Reads were trimmed to 120bps and the first 10bps were trimmed from each read. Reads were mapped to the *P. picta* reference genome using hisat2 v2.1.^84^ Only uniquely mapped reads that aligned as pairs were retained for further analysis. Average mapping efficiency was 97.7% with an average mapped library size of 67,004,220 reads (Supplemental Table 2). Alignment bam files were sorted and indexed using samtools v1.3.1.^85^ Aligned reads were assigned to genic regions using featureCounts v2.0.1.^86^

Gene expression analysis was conducted in R v4.1 using edgeR v3.34.1.^87^ Genes with zero counts in all samples were removed. We then filtered the data to retain only genes with >10 reads in each female or male sample. This filtered dataset was then used to calculate normalization factors using the RLE method in the calcNormFactors() function. We estimated robust dispersions using the estimateDisp() followed by the estimateGLMRobustDisp() function. Normalized counts per million (cpm) values were then extracted from the data to compare gene expression levels between males and females and between genes located on the X chromosome compared to the autosomes. For the quantile expression analysis, the quantile() function from the R stats package was used to bin genes into the 0-25, 25-50, 50-75, and 75-100th percentiles based on gene expression values in males.^31^ A Wilcoxon rank sum test was used to identify differences in gene expression levels between groups using an FDR corrected p-value <0.05.

### Bisulfite Sequencing sampling and analysis

Muscle, liver, and gonad tissues were sampled from adult male and virgin female *P. picta* as described above. A total of 15 males and 15 females were sampled. To minimize the effect of genetic differences and developmental factors contributing to inter-individual variation in DNA methylation patterns,^88^ we pooled tissues from five male or five female tissue samples to generate a total of three male and three female non-overlapping sample pools for DNA isolation and sequencing with the exception of ovary tissue where we were only able to extract enough DNA from 2 of the 3 replicates. DNA was purified from pooled tissue samples using Qiagen DNeasy spin columns with on-column RNase A treatment following the manufacturer’s protocol. Purified genomic DNA samples were frozen at −20 °C, and then shipped overnight on dry ice to the McGill University and Génome Québec Innovation Centre for bisulfite conversion, NEBNext^®^ Enzymatic Methyl-seq library preparation, and sequencing on the Illumina 6000 PE150 platform.

WGBS sequencing reads were quality filtered and adapters were trimmed using trimmomatic v0.36.^79^ Reads were scanned with a 4-base sliding window and discarded when the average phred score was <15. Leading/trailing bases with a phred score <3 were also removed. Trimmed and filtered reads were mapped to the *P. picta* reference genome using BSBolt.^89^ Only uniquely mapped reads that mapped as pairs were retained for further analysis. The average mapping efficiency was 86.5 % and the average library size is 105,671,545 reads (Supplemental Table 5). BAM alignment files were prepared for duplicate removal using samtools v1.15.1 fixmate followed by samtools sort.^85^ Duplicate reads were removed using samtools markdup and then indexed. We then used the BSBolt CallMethylation function to calculate DNA methylation levels at single nucleotide resolution. Differential methylation analysis was conducted using the R package DSS v2.4.^90^ Differentially methylated loci (DMLs) and differentially methylated regions (DMRs) between males and females were identified for each tissue using the dmlTest() function with smoothing = TRUE followed by callDML() and callDMR() respectively. CpG islands were identified using a hidden markov model using the makeCGI_1.3.4 R package.^91^

### Motif enrichment and YY1 analysis

Motif enrichment analysis for differentially methylated regions on the X chromosome was conducted using findMotifsGenome.pl in Homer v4.11.1.^92^ Sequences from DMRs on the X chromosome were used as the query sequences and sequences from DMRs on the autosomes were used as background sequences. We used the matchPattern() function in the Biostrings v2.60.2 R package to search for YY1 motifs in the *P. picta* genome and to also identify repetitive regions containing YY1 motifs. DNA methylation of YY1 motifs was calculated as the mean DNA methylation value of CpG loci +/- 500bps from a YY1 motif.

### De novo repeat element analysis

For the de novo identification of repetitive elements in the *P. picta* genome we used the RepeatModeler v2.0.1^93^ to generate a repeat element library. We then used RepeatMasker V4.1.2^94^ to annotate the repeat elements in the *P. picta* genome and to calculate kimura divergence estimates. Repetitive elements containing YY1 motifs were aligned using Muscle v.5.1.^95^ The output file from the Muscle alignment was then processed using FastTree V2.1.^96^ The phylogeny from FastTree was visualized using the ggtree v3.0.4 package in R. To compare TE length and abundance between chromosomes, we calculated the total length (bp) and number of TEs in 200kb non-overlapping sliding windows. The mean total length and count per 200kb for each scaffold were then plotted against each other in a scatter plot with standard error bars and a linear regression line.

## Data Availability

The *Poecilia picta* genome assembly is available from the NCBI Genome database BioProject PRJNA862953. Raw RNAseq sequence files are available from the NCBI sequence read archive (SRA) BioProject PRJNA884377 (SRA accession #’s SAMN31026551 - SAMN31026568). Raw whole genome bisulfite sequence files are available from the SRA Bioproject PRJNA884372 (SRA accession #’s SAMN31025781 - SAMN31025797). Computer code used for data processing and analysis are available at https://github.com/manklab/Metzger_et_al_picta_DC.

## Acknowledgements

We would like to thank Jacelyn Shu for the scientific illustrations in Figure 4, for help with animal husbandry, and tissue dissections for karyotype analysis. We thank Clara Lacy for the drawings of male and female *Poecilia picta*. We would also like to thank Wouter van der Bijl for assistance with R and members of the Mank lab for helpful and constructive suggestions on this manuscript. This work was funded by grants from the European Research Council (grant number 680951), NSERC and CFI, as well as a Canada 150 Research Chair to JEM. We also acknowledge funding from a Dovetail Genomics Tree of Life Award to DCHM and funding from NSERC Undergraduate Student Research Award grant awarded to IP.

## Author Contributions

DCHM and JEM conceived the study and wrote the manuscript. DCHM performed tissue collection and sample preparation for genome sequencing, WGBS, and RNAseq data, performed bioinformatic analysis of WGBS and RNAseq data, motif enrichment, and characterization of YY1 repetitive elements. IP identified and characterized repetitive elements in the *P. picta* genome. BM and AA performed the karyotype analysis. BAS sequenced the mitochondrial genome, performed YY1 amino acid alignments, and helped with sample collection. LJMV helped with sample collection.

---

**Supplemental Figure 1.**
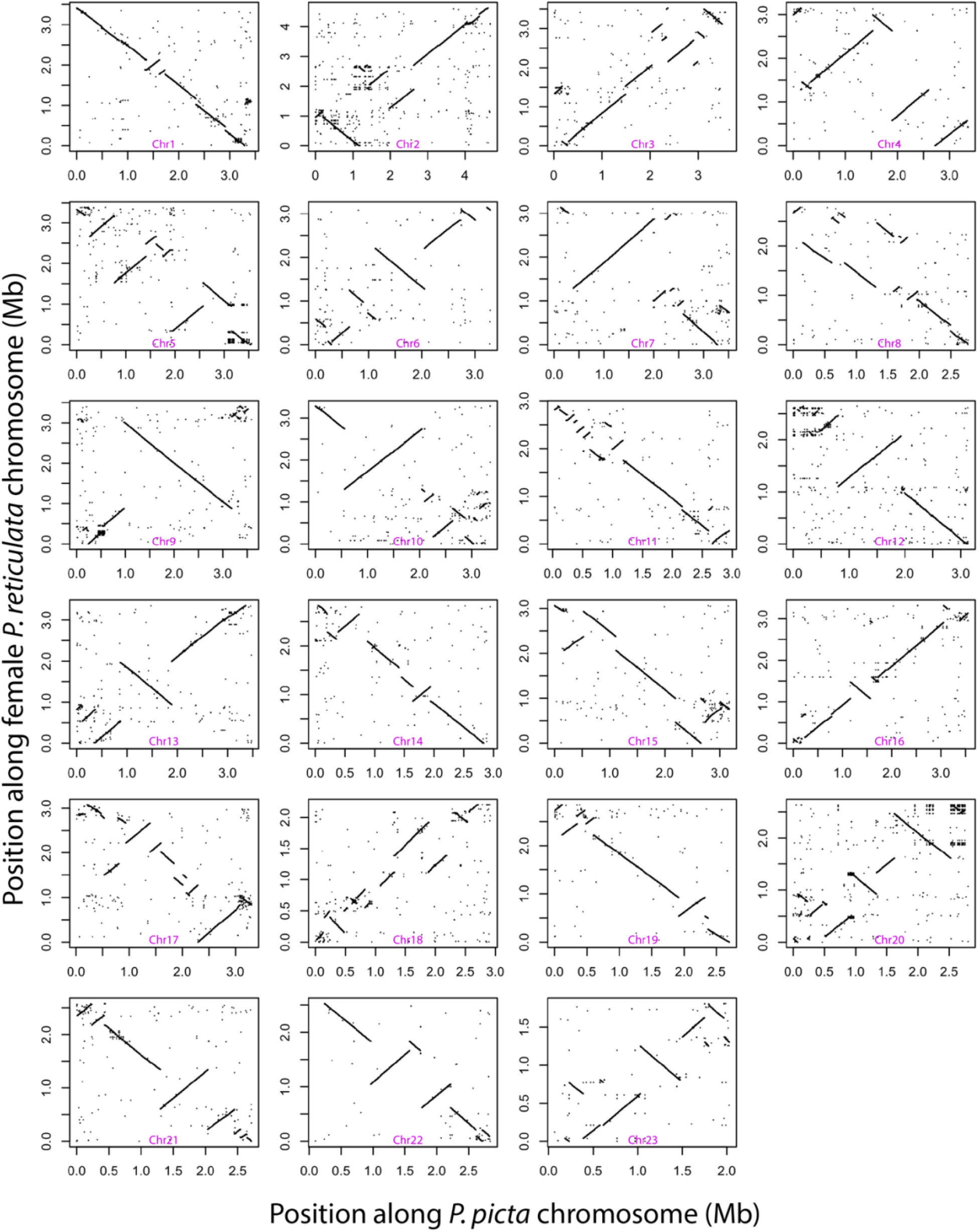
Large-scale rearrangements between our *P. picta* and female *P. reticulata* genome assemblies. The X chromosome is Chrl2.

**Supplemental Figure 2.**
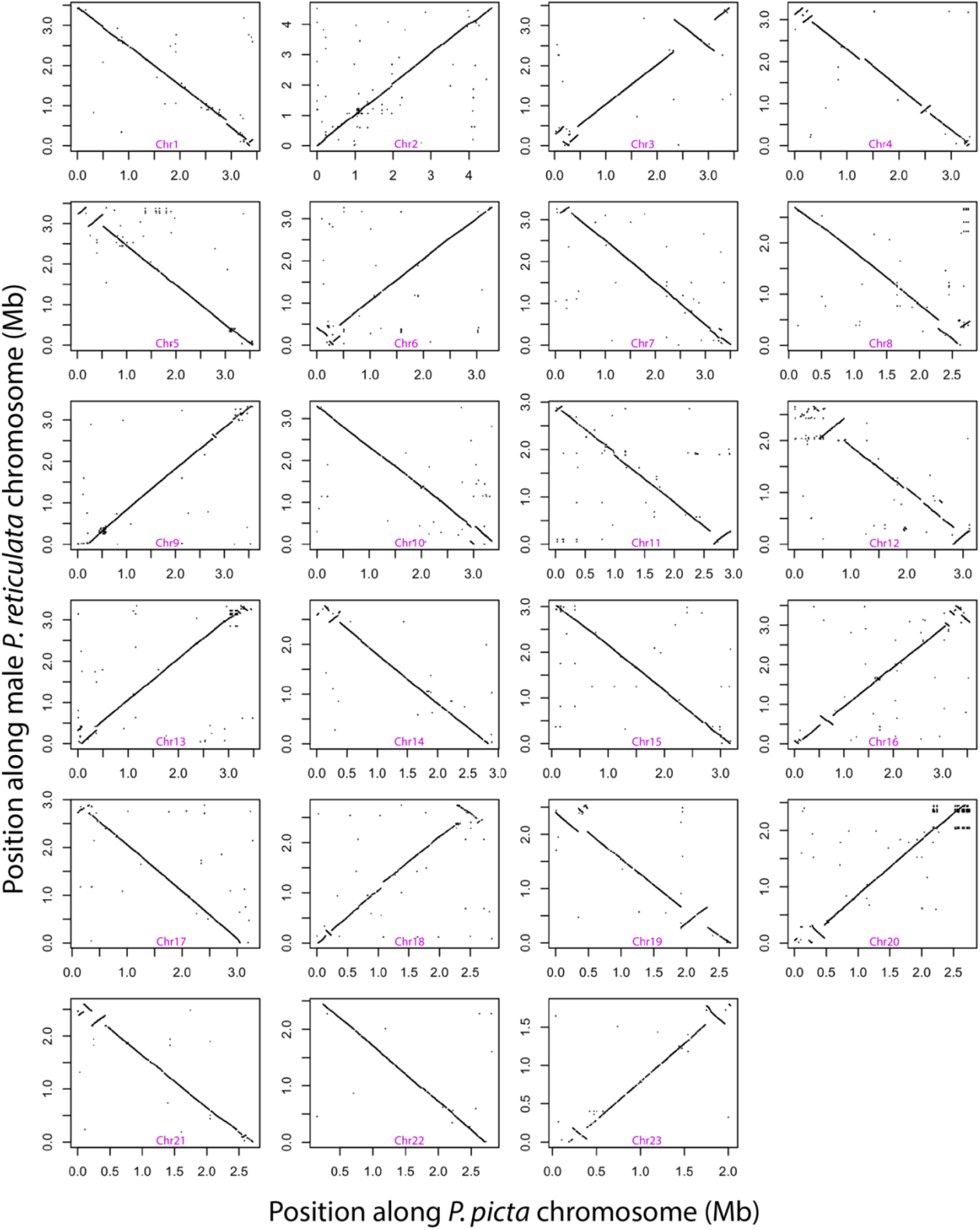
Substantial synteny between our *P. picta* and the male *P. reticulata* genome assemblies. The X chromosome is Chrl2.

**Supplemental Figure3:**
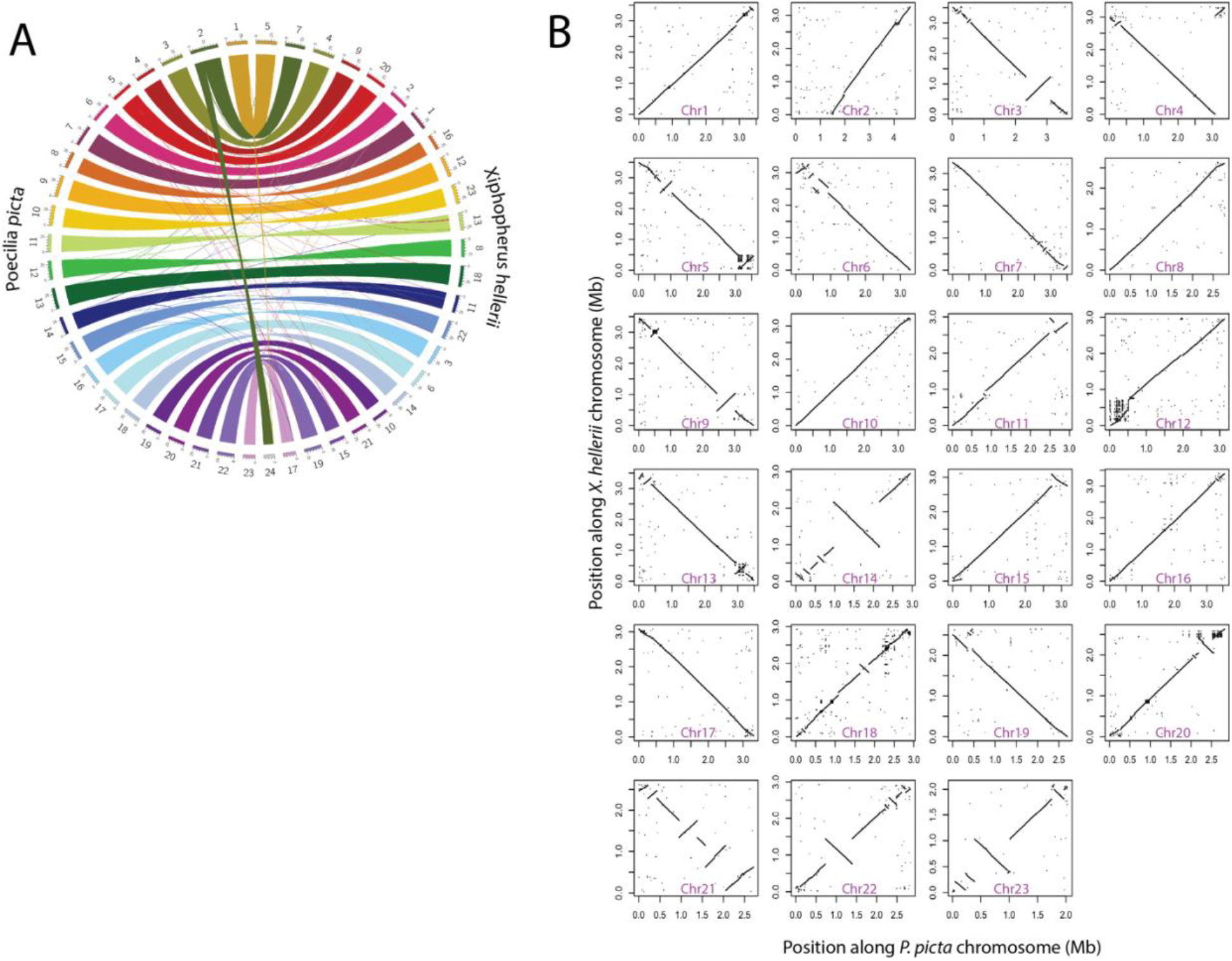
Sequence comparison between our *P. picta* genome and the *X. hellerii* genome. A) Sequence synteny between the female *P. picta* and the female *P. helleri* genomes presented as a Circos plot. Chromosomes are labeled 1-23 and are differentiated by colour around the outer ring with *P. picta* chromosomes on the left and *P. helleri* chromosomes on the right (numbered 1-24). The colour of the connecting lines indicates the chromosomal origin of the sequence from the *P. picta* genome. B) Synteny scatter plot between *P. picta* and *X. helleri* chromosomes. Each plot is a separate chromosome with the *P. picta* chromosome indicated on each plot. The *P. picta* X chromosome is chr12.

**Supplemental Figure 4.**
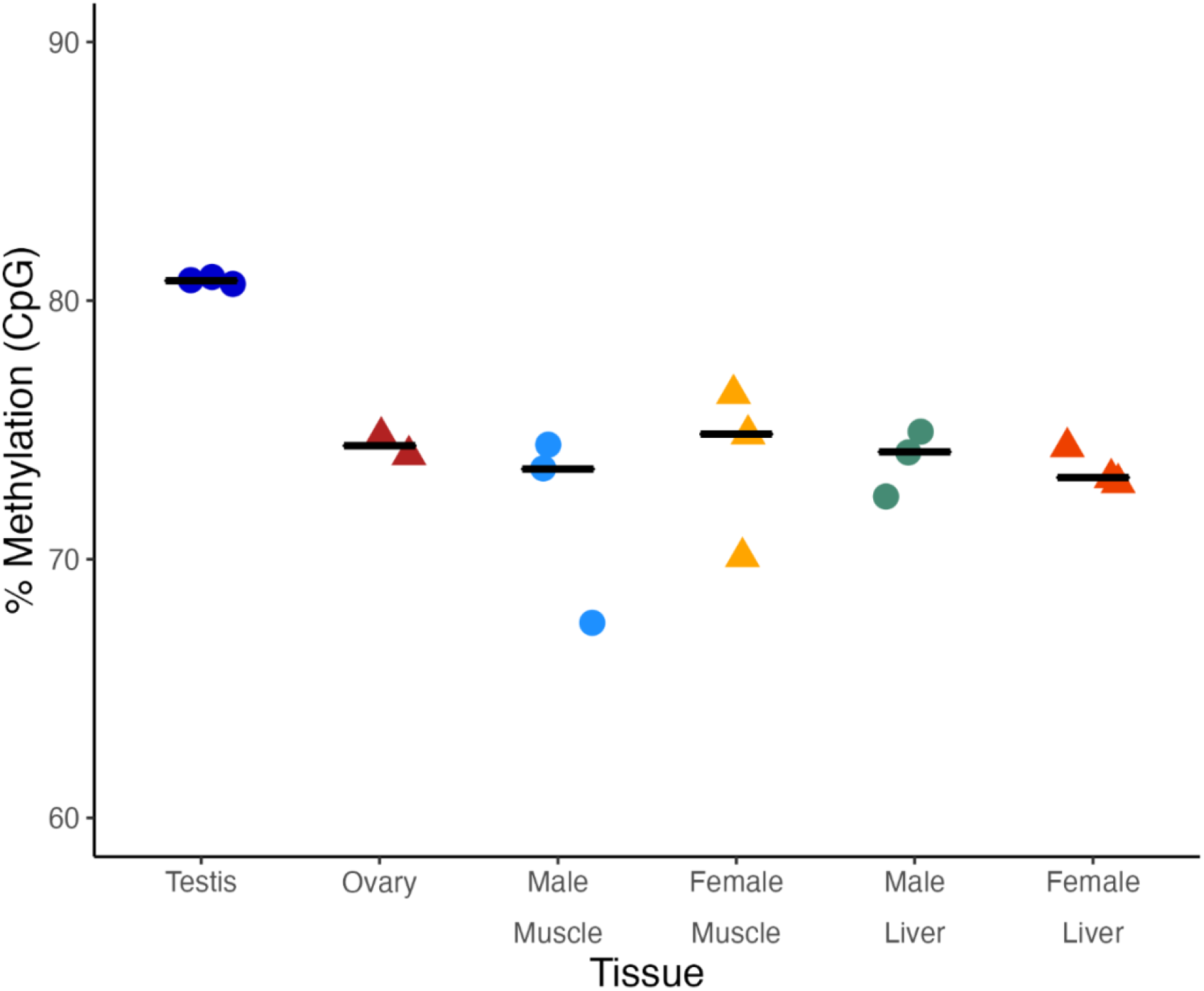
Methylation varies by tissue and sex. Whole genome DNA methylation levels for *P. picta* testis (T), ovary (O), male muscle (MM), female muscle (FM), male liver (ML, and female liver (FL) tissue in P. picta. Circles represent individual data points from male tissue. Triangles represent individual datapoints from female tissue. Percent DNA methylation values were calcualted from whole genome bisulfite sequencing data for sites with >10 reads.

**Supplemental Figure 5.**
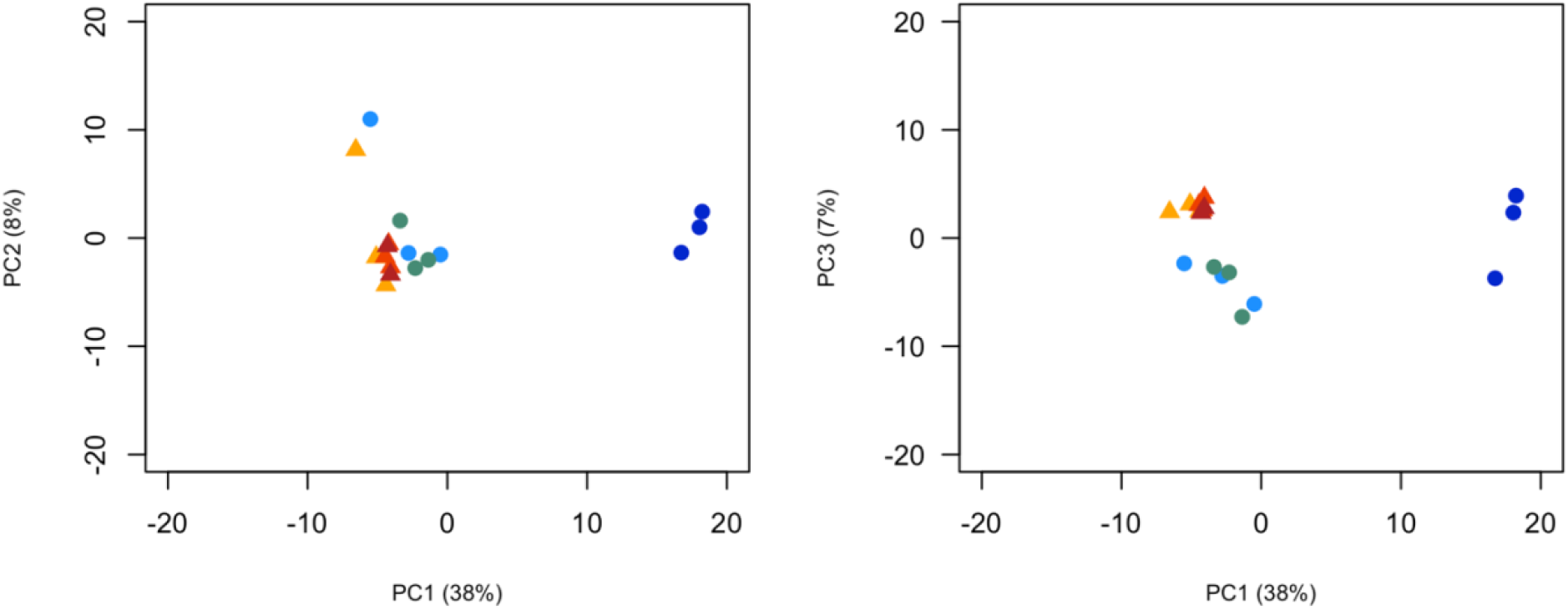
Testis is a methylation outlier. Principle componenet analysis of *P. picta* DNA methylation data calculated from whole genome bisulfite sequencing data. Circles represent samples from male tissue. Triangles represent samples from female tissue. Colors are the same as in Supplemental Figure 4.

**Supplemental Figure 6.**
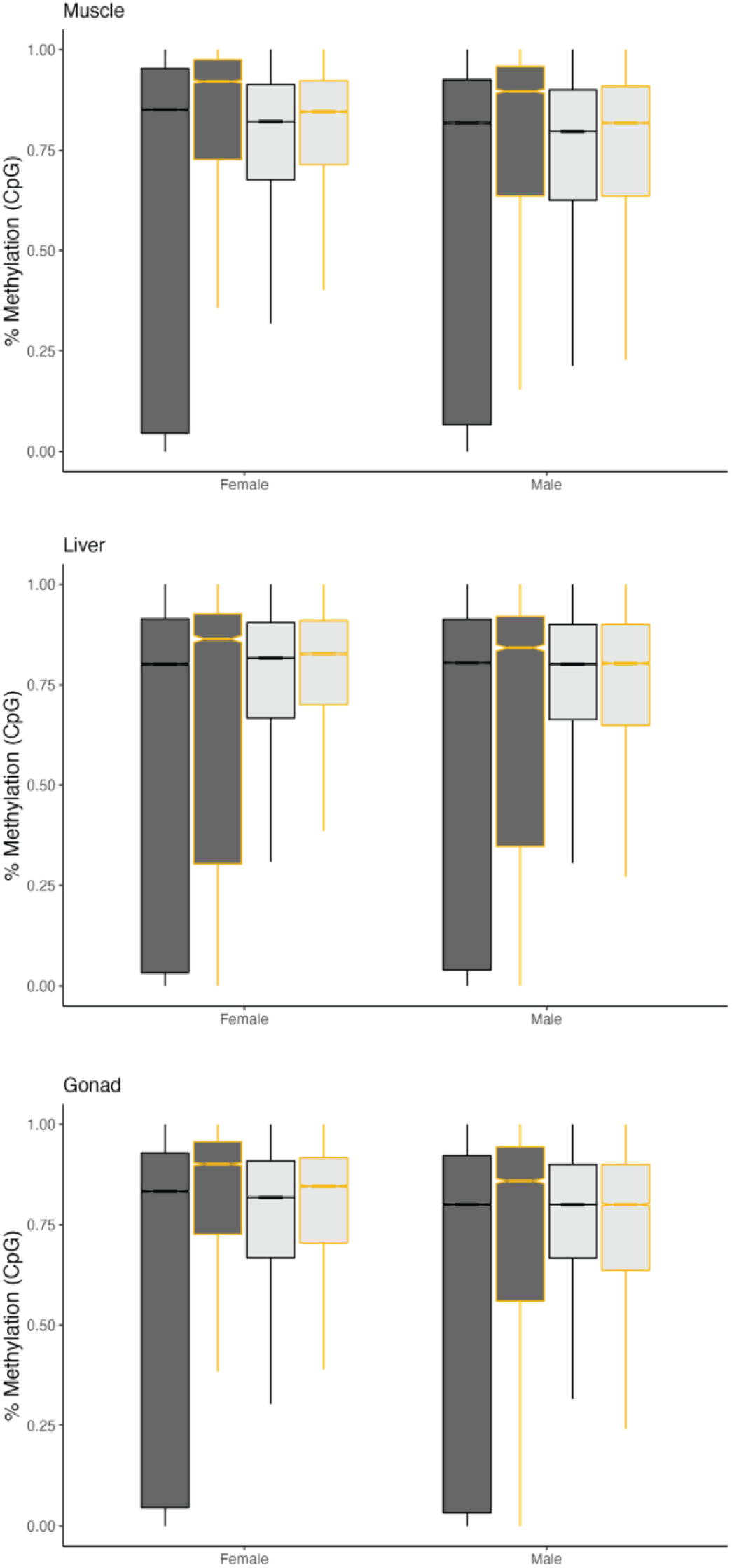
CpG islands on the X chromosome are more highly methylated compared to autosomes in both males and females. *P. picta* DNA methylation values of individual CpG located in CpG islands (dark grey) or outside of CpG islands (light grey) on the sex chromosome (orange outline) or autosomes (black outline).

**Supplemental Figure 7.**
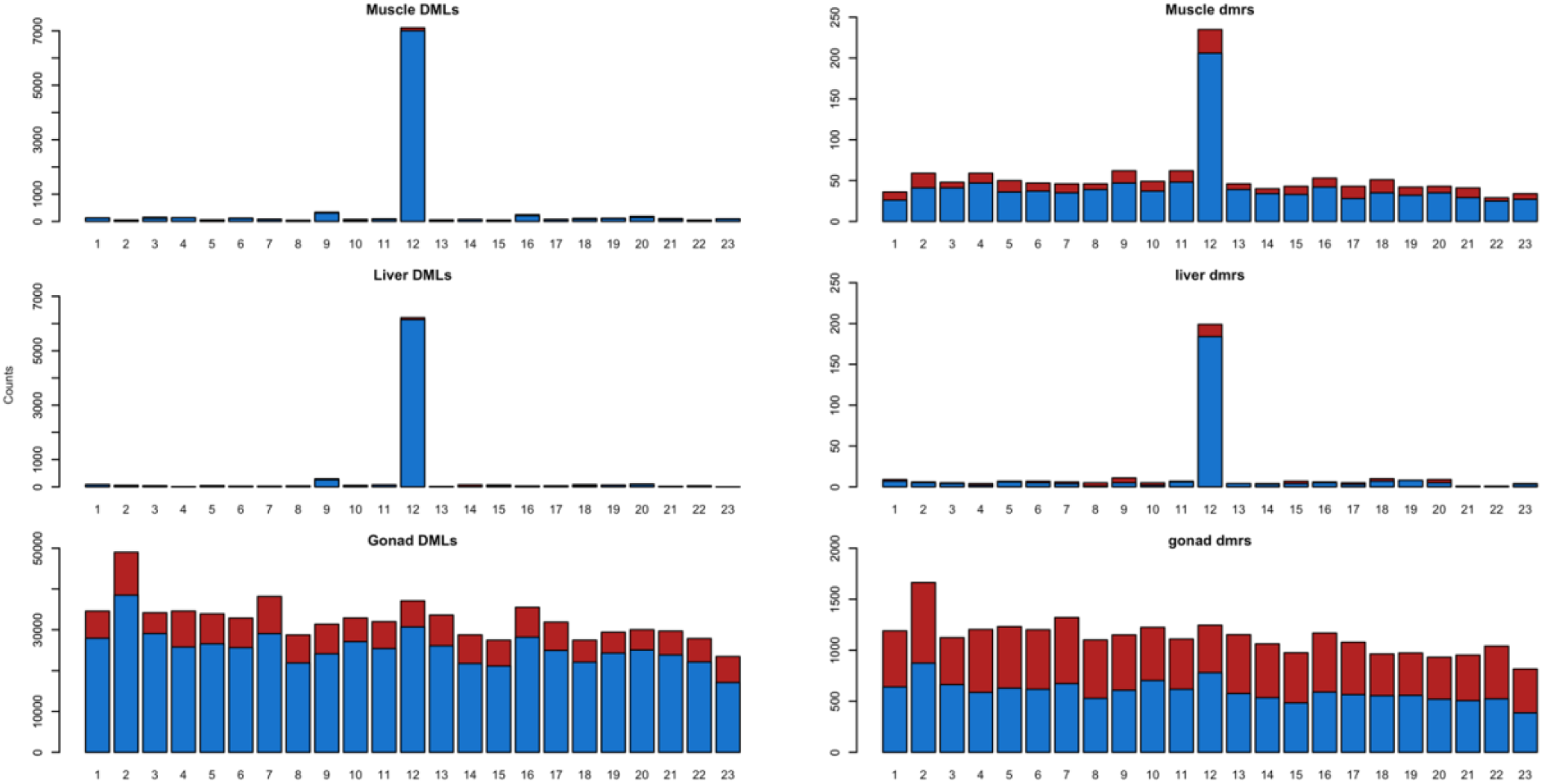
The X chromosome is hypomethylated in *P. picta* males. Chromosomal distribution of differntially methylated loci (DMLs) and differentially methylated regions (DMRs) in muscle, liver, and gonad tissue presented as stacked bar plots where the height is the total number of DMLs or DMRs identified on an individual chromosome. Red shaded regions are DMLs or DMRs that are hypomethylated in female tissue Blue shaded regions are DMLS or DMRs that are hypomethylated in male tissue.

**Supplemental Figure 8.**
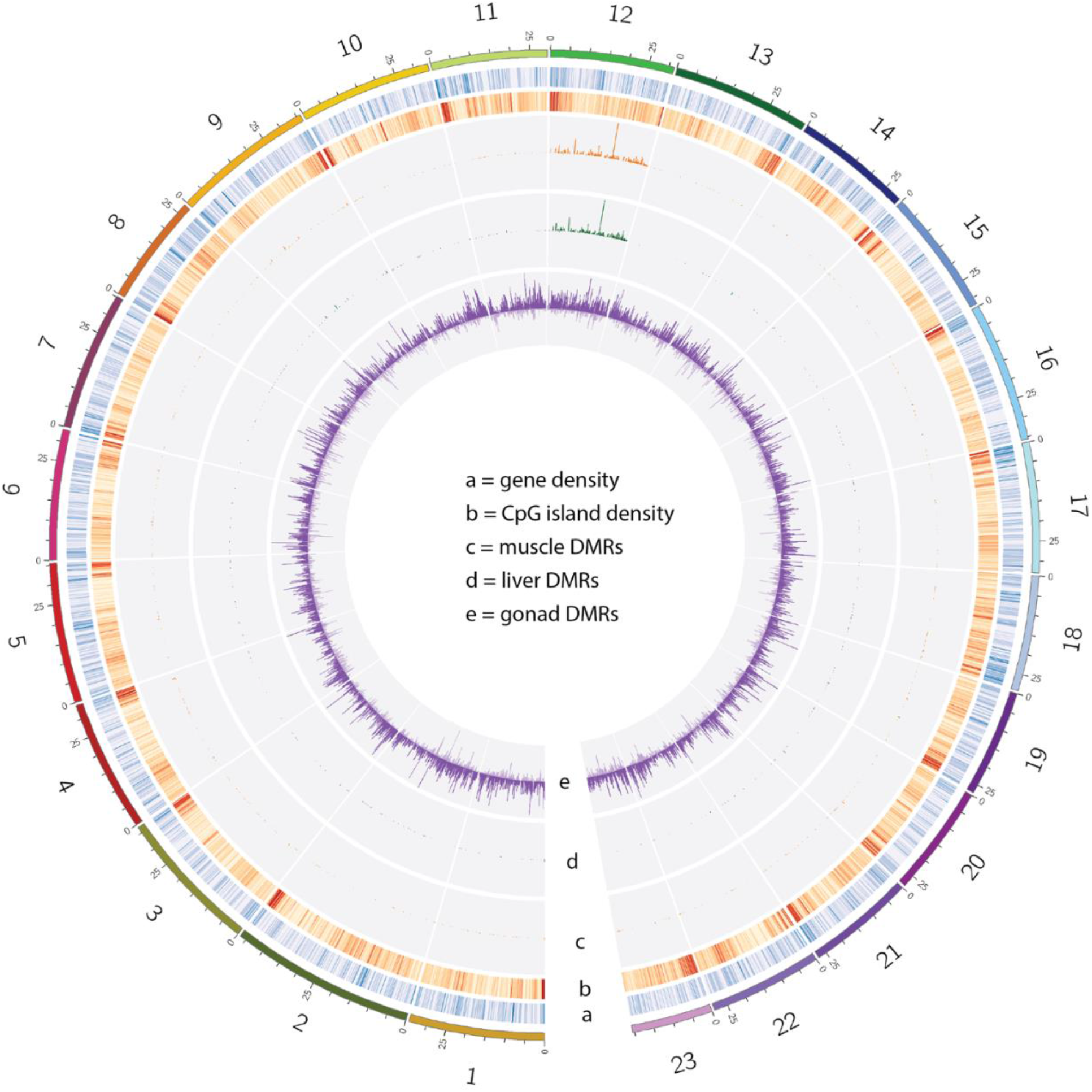
Chromosome 12, the X chromosome, is hypomethylated in *P. picta* male somatic tissues. Circos plot depicting sex-specifc DNA methylation patterns in *P. picta*. Outer ring indicates the chromosome number and position in megabases. Chromosome colours are the same as in Figure 1. Tracks a & b are heatmaps depicting gene (blue) and CpG island (Red) densities respectively, in 100kb bins. Darker colors in the heatmap depict higher counts. Tracks c-d depict histograms of the number of differentially methylated CpG loci (DMLs) in 100kb bins in muscle (orange), liver (green), and gonad (purple) tissues respectively. Histogram bars extending outward represent DMLs that are hypomethylated in males while bars that extend towards the center of the plot represent DMLs that are hypomethylated in females.

**Supplemental Figure 9.**
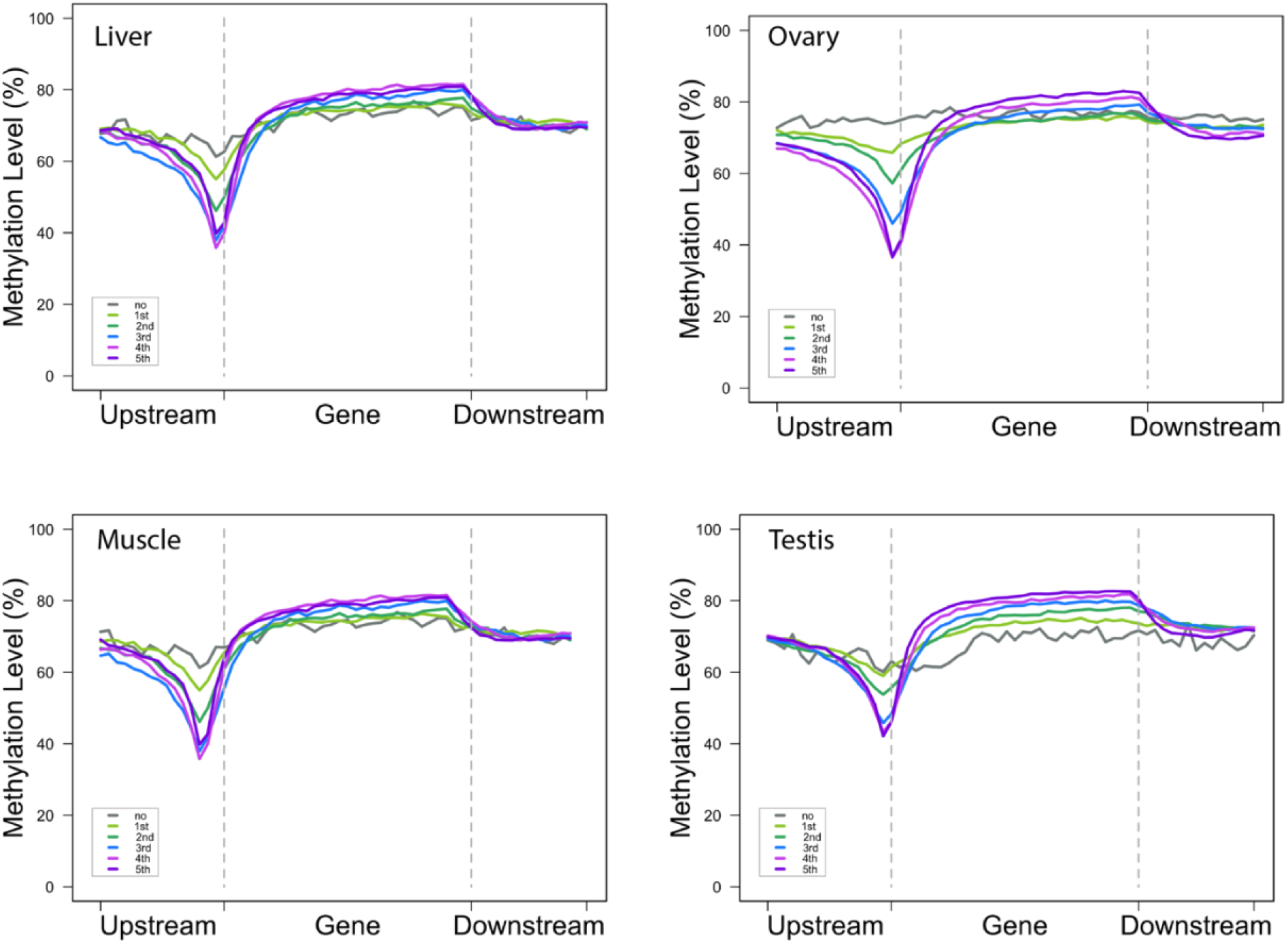
Association of DNA methylation and genes ranked by gene expression level in *P. picta*. DNA methylation values for genes with no expression are in gray, genes with low expression are in green, medium expression in blue and pink, and the highest expression in purple.

**Supplemental Figure 10.**
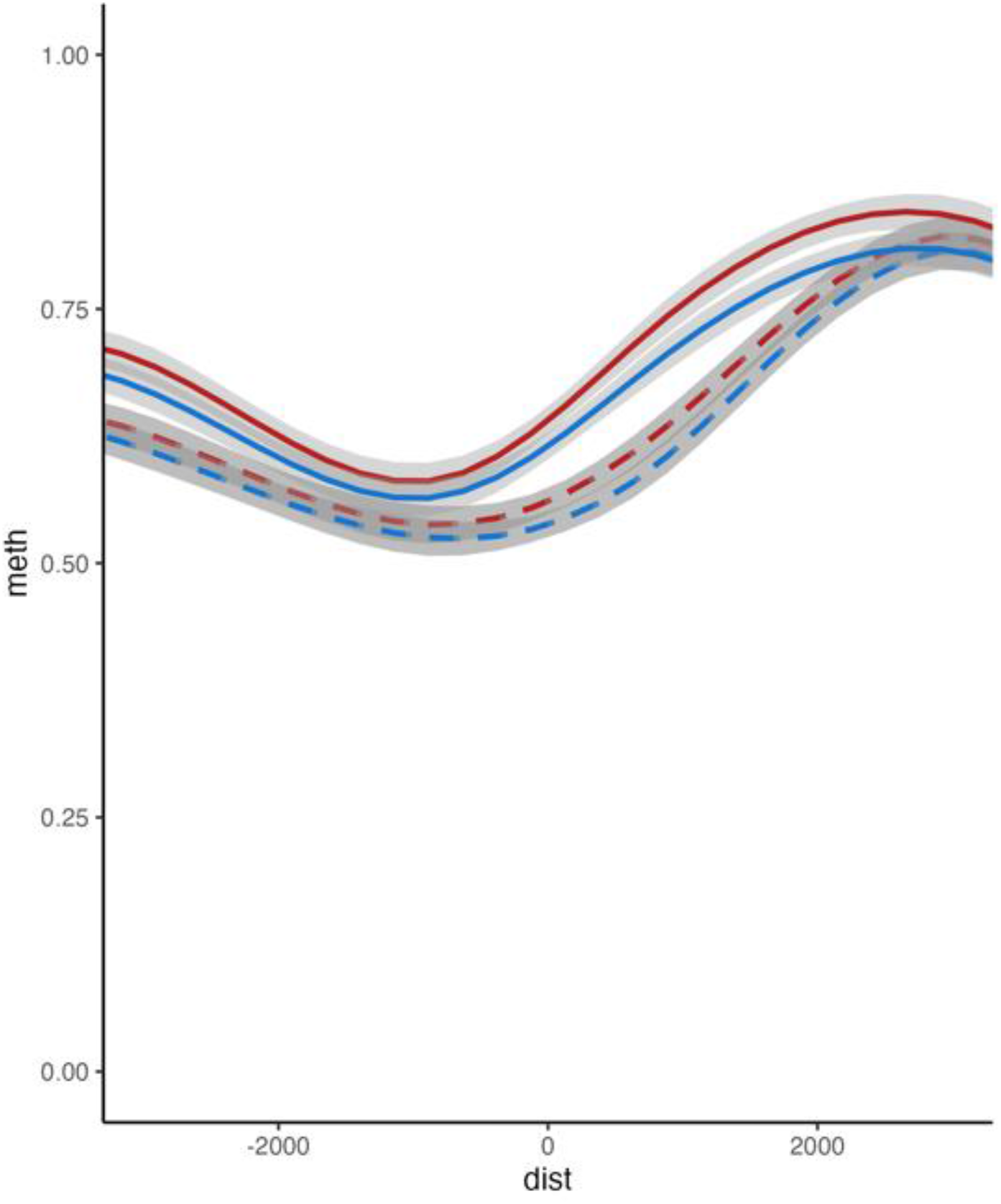
CpG islands in gene promoters on the X do not exhibit sex-biased differential methylation. DNA methylation values in the promoter region of genes on the X chromosome that contain CpG islands for female (red) and male (blue) muscle (solid line), liver (dashed line), and gonad (dotted line). Shaded grey region denotes 95% confidence intervals.

**Supplemental Figure 11.**
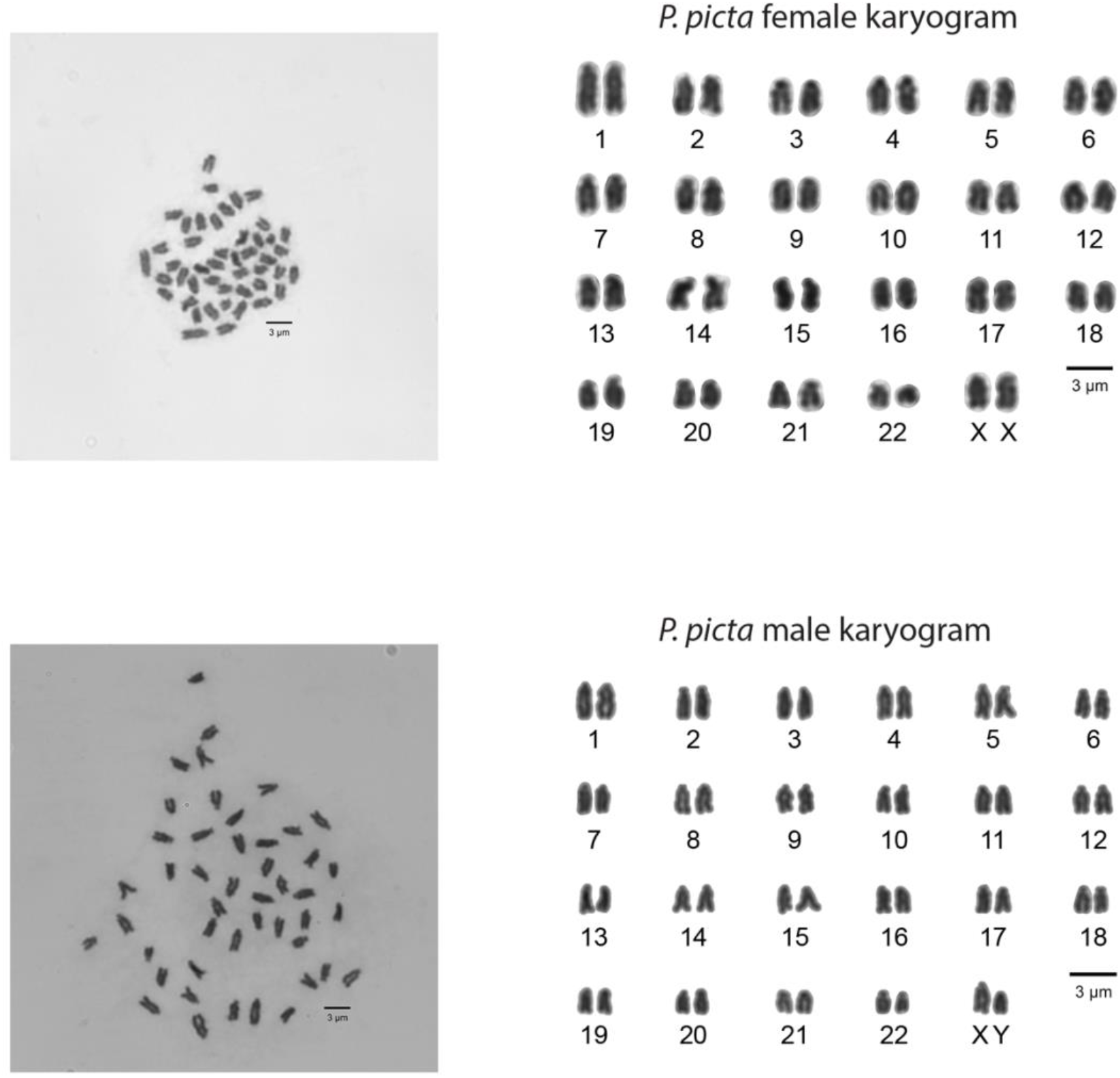
Male and female X chromosome appear similar in size. Karyograms from male and female *P. picta* chromosome spreads from our laboratory population originally sourced from Suriname. The Y chromsome appears degenerated compared to the X, while the X chromsomes in males and females are similar in size at this resolution. The left panel shows the chromosome spreads from which the karyograms were sourced.

**Supplemental Figure 12.**
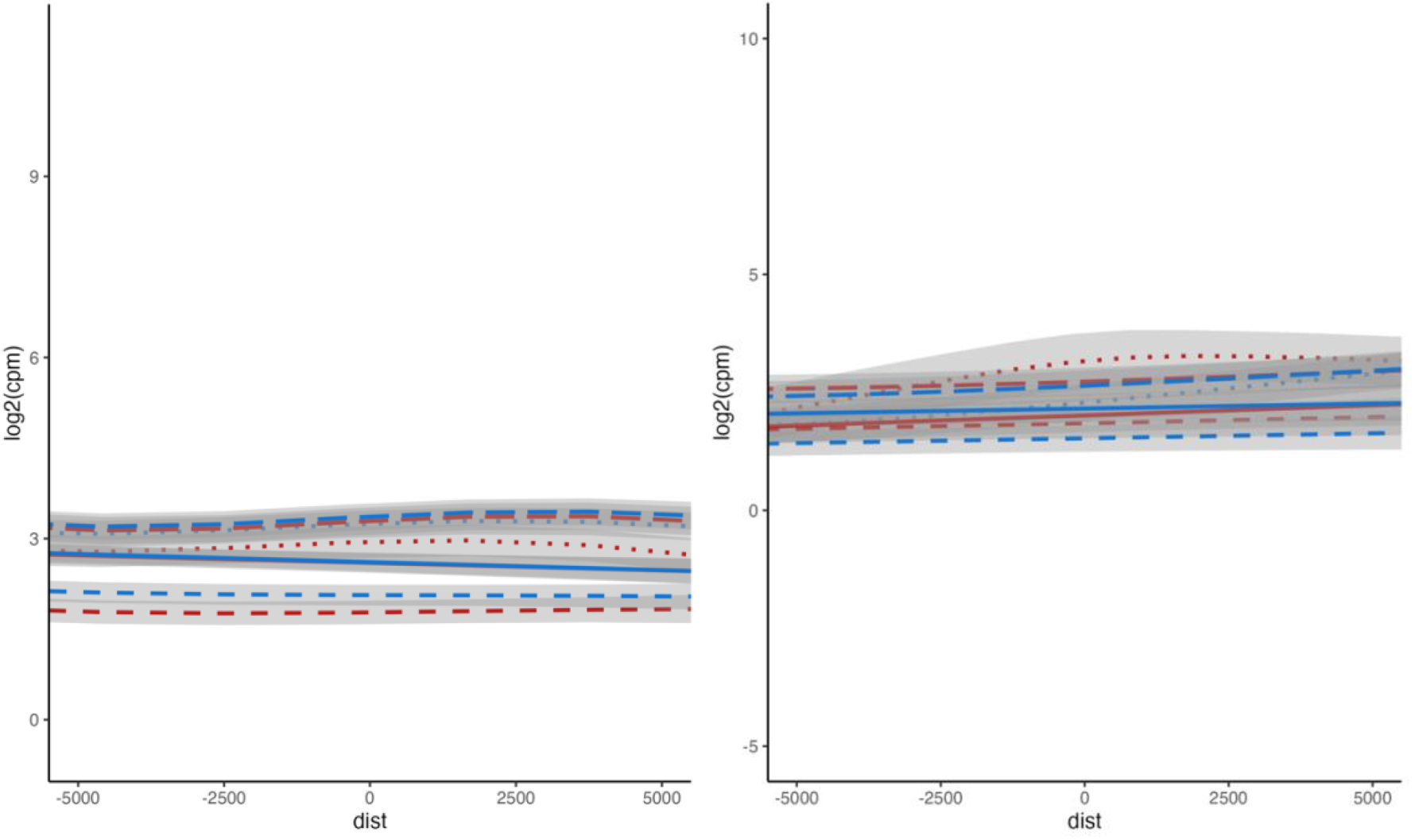
Proximity to YY1 motifs is not associated with sex-biased gene expression. Gene expression values for genes 5kb upstream and 5kb downstream of YY1 motifs on the sex chromosome (left panel) or on autosomes (right panel) in female (red) and male (blue) muscle (solid line), liver (dashed line), and gonad tissue (dotted line). Shaded grey areas represent 95% confidence intervals.

**Supplemental Figure 13.**
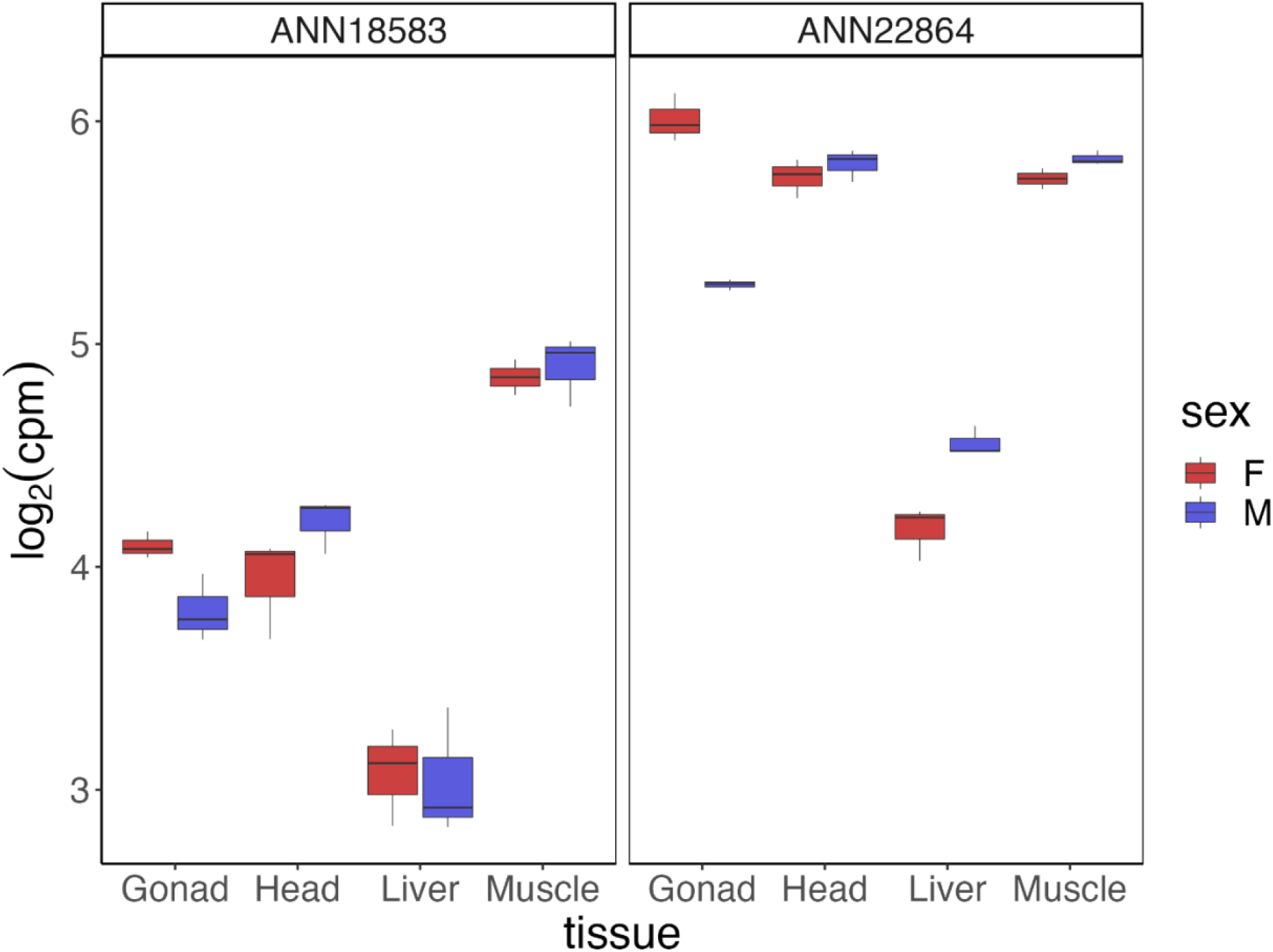
YY1 orthologs are ubiquitously expressed in head, muscle, liver, and gonad tissues. Gene expression values for two YY1orthologs (ANN18583 and ANN22864) in the *P. picta* genome for female (F, red), and male (M, blue) gonad, head, liver, and muscle tissue. Data is present as log2 counts per million normalized for library size as box and whisker plots. The horizontal line of the box and whisker plot is the median, the box denotes the 25th and 75th percentile, and “whiskers” are 1.5 times the interquartile range.

**Supplemental Figure 14.**
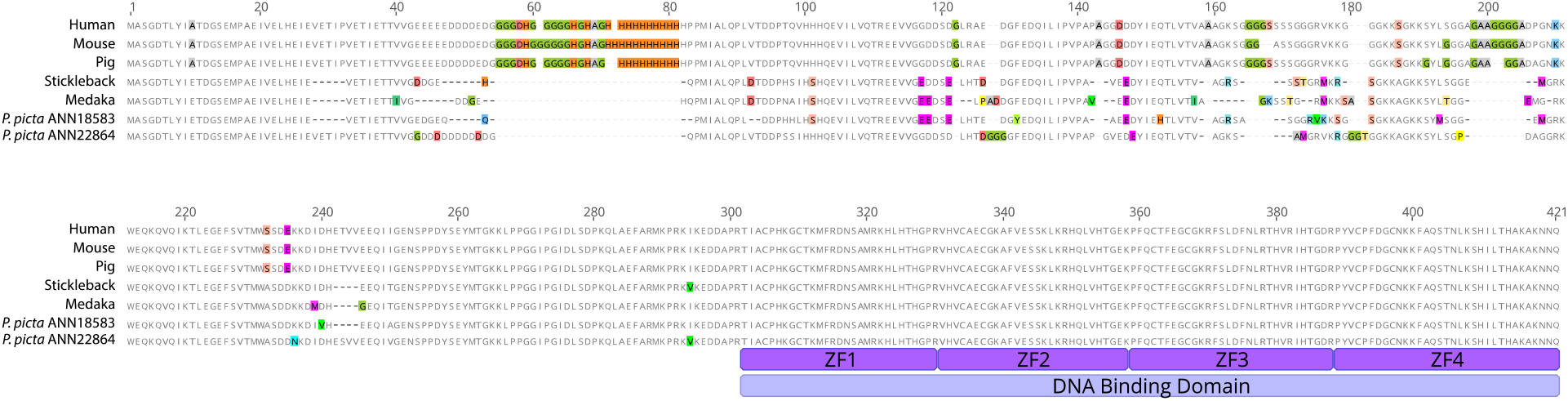
The DNA binding domain of the YY1 protein is conserved between mammals and fishes. Alignment of amino acid sequecnes from human, mouse, pig, stickleback, medaka, and two isoforms from *P. picta*. Coloured amino acid residues depict mismatches between sequences. Purple ZF blocks indicate the four C2H2-type zinc finger domains of the YY1 DNA binding domain.

**Supplemental Figure 15.**
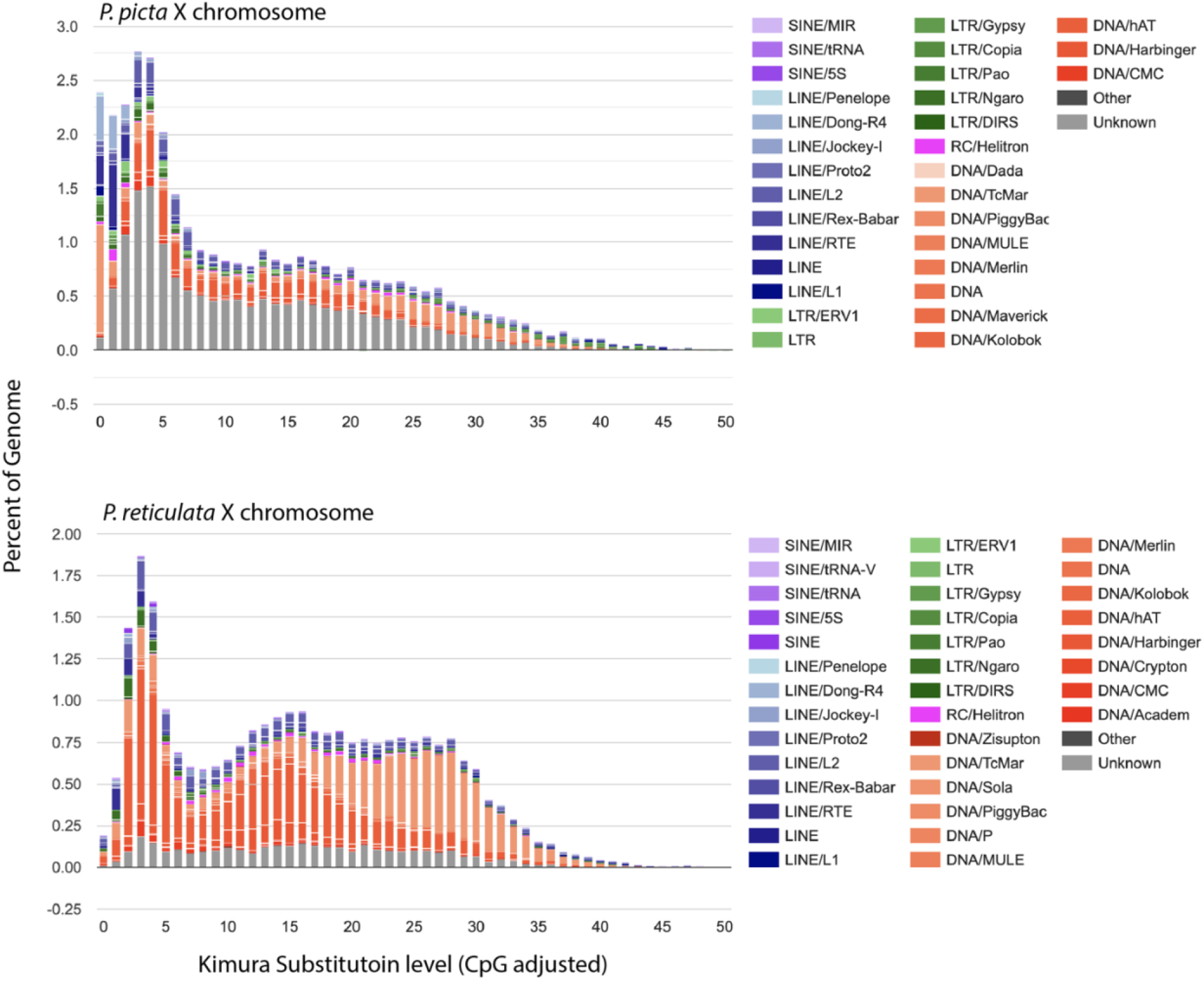
The *P. picta* genome contains a group of recently derrived repetative elements compared to *P. reticulata*. Kimura substitution values used as a proxy for the expansion history of repretitive elements for the *P. picta* (upper panel) and *P. reticulata* (lower panel) X chromosomes. Bargraph depicts the proportion of the X chromosome that contains repetitive elements for a given kimura substitution level. Colors indicate RE annotations. The older the repetitive element expansion the higher the Kimura substitution level indicating more sequence divergence from the consensus.

**Supplemental Table 1.** *P. picta* genome assembly statistics.

|  |  |
| --- | --- |
| HiRise assembly size (bp) | 743,972,568 |
| Number of Scaffolds | 301 |
| N50 | 33,053,486 |
| L50 | 11 |
| N90 | 27,142,806 |
| L90 | 21 |
| Largest scaffold | 46,707,787 |
| Genome completeness (BUSCO) | 97.65 % |
| Total number of genes | 27,764 |
| Total coding region | 46,815,666 |
| Repeat % | 29.55 % |

**Supplemental Table 2.** Comparison of average expression between the sex chromosome and autosomes across tissues and sex.

| Tissue | X | Y | p-value* |
| --- | --- | --- | --- |
| Gonad | Male X genes | Male Autosomal genes | 4.915e-12 |
|  |  | Female X genes | 6.443e-07 |
|  |  | Female Autosomal genes | 1.036e-07 |
| Muscle | Male X genes | Male Autosomal genes | 0.6724 |
|  |  | Female X genes | 0.7923 |
|  |  | Female Autosomal genes | 0.2407 |
| Liver | Male X genes | Male Autosomal genes | 0.9855 |
|  |  | Female X genes | 0.6603 |
|  |  | Female Autosomal genes | 0.1029 |
| Head | Male X genes | Male Autosomal genes | 0.94 |
|  |  | Female X genes | 0.3903 |
|  |  | Female Autosomal genes | 0.8463 |

\*p-values calculated using Wilcoxon rank sum test between data in column X and data in column Y

**Supplemental Table 3.**
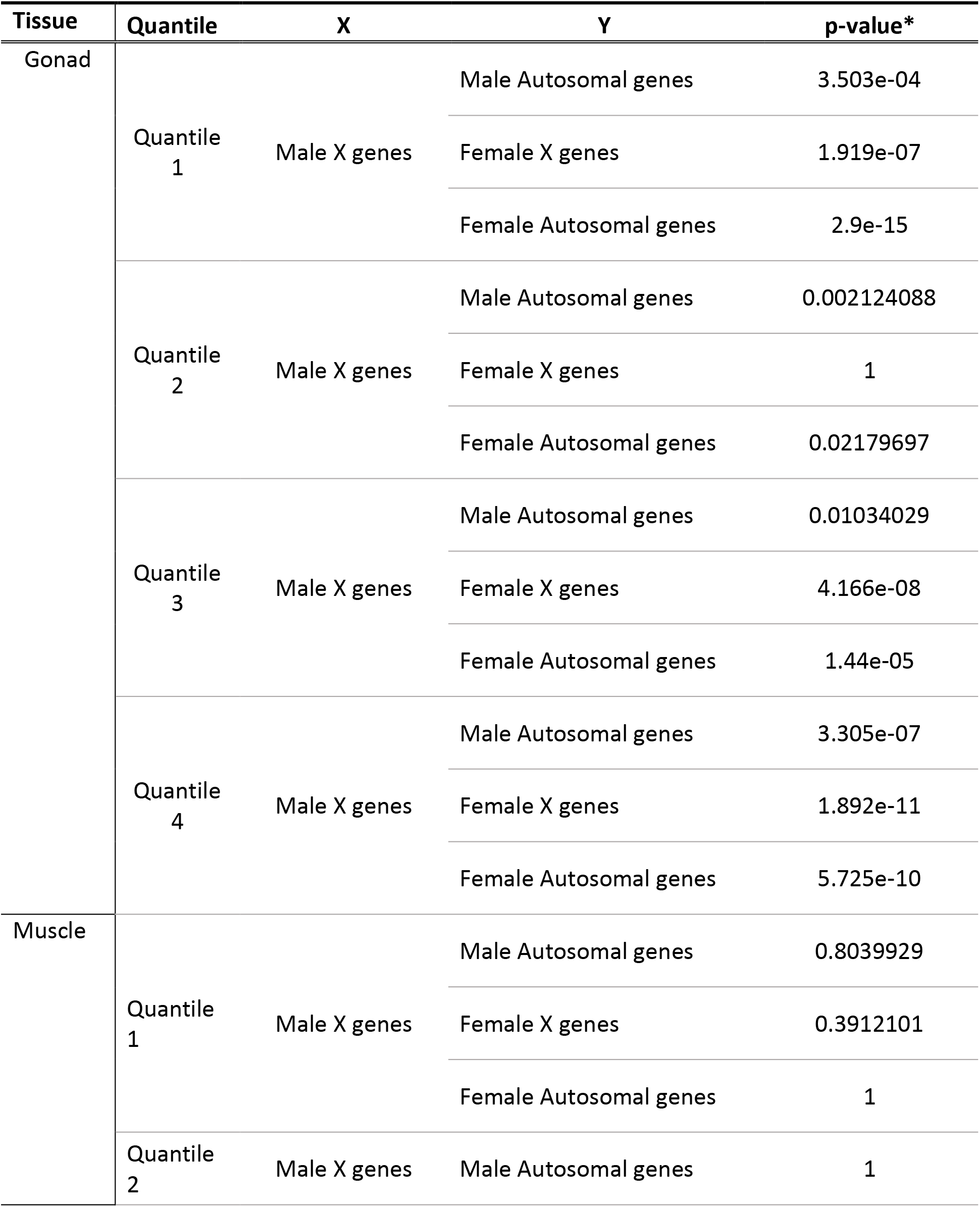
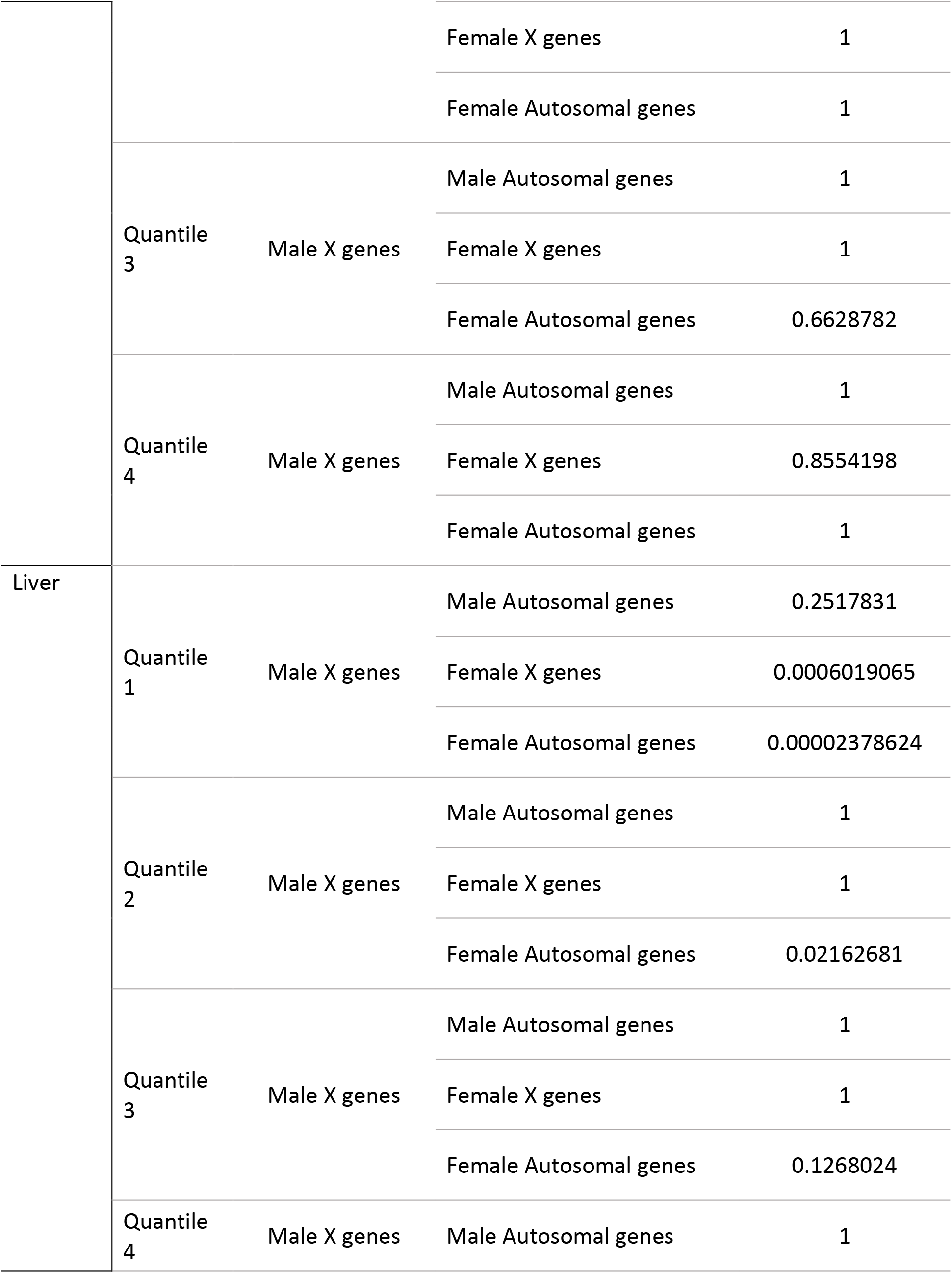
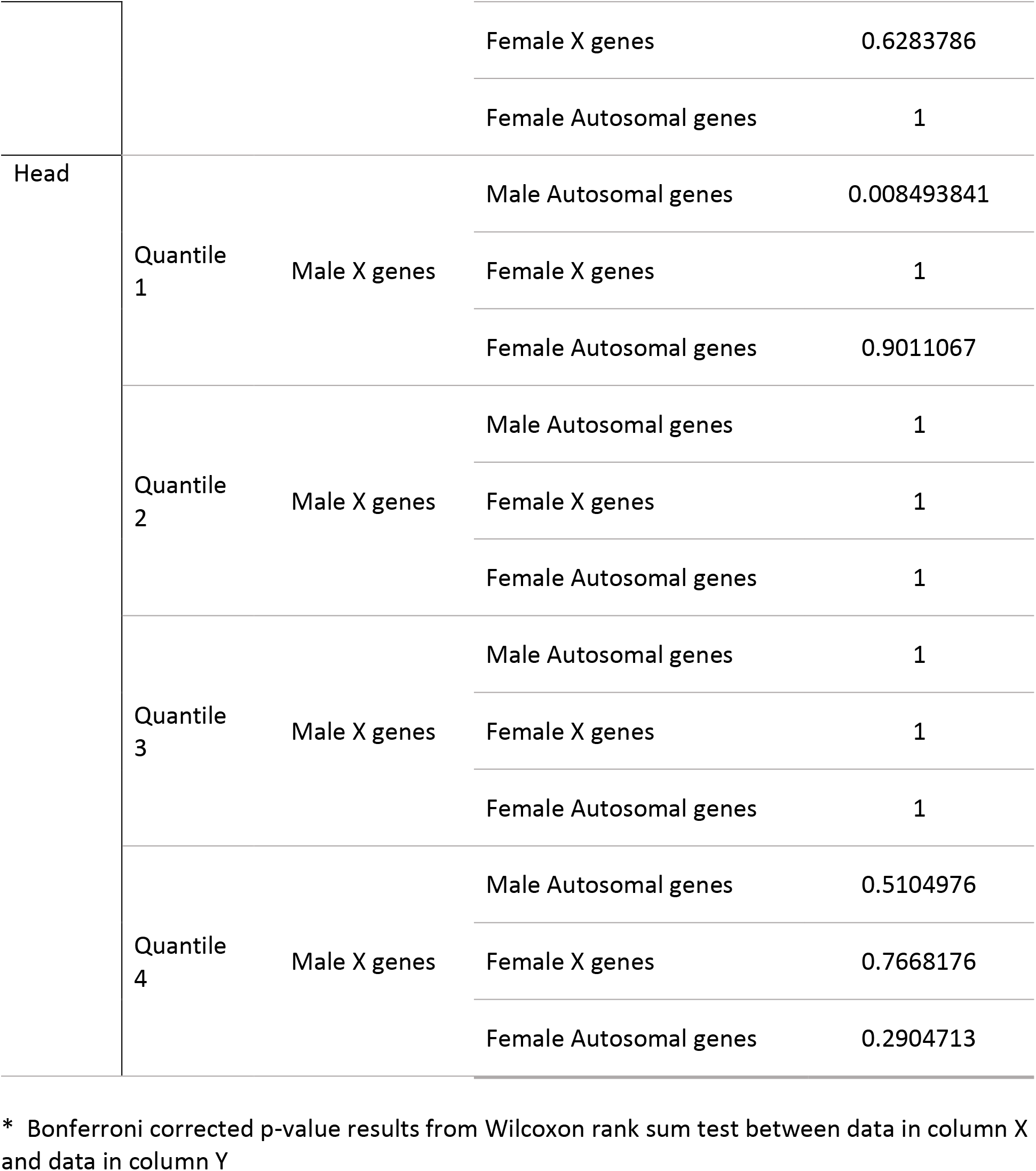
Wilcoxon rank sum test statistics for dosage compensation as a function of expression level.

**Supplemental Table 4.** RNA-Seq data for *P. picta*.

| Tissue | Sex | Biological Replicate | # reads | # reads mapped as pairs | % reads mapped |
| --- | --- | --- | --- | --- | --- |
| Muscle | Male | 1 | 144636107 | 118061726 | 88.28% |
|  |  | 2 | 169476423 | 138122808 | 89.86% |
|  |  | 3 | 194995685 | 158505438 | 90.33% |
|  | Female | 1 | 148583536 | 120583282 | 91.11% |
|  |  | 2 | 165442661 | 130740620 | 89.83% |
| Liver | Male | 1 | 42653690 | 42602048 | 99.88% |
|  |  | 2 | 46358957 | 46311114 | 99.90% |
|  |  | 3 | 42446289 | 42401986 | 99.90% |
|  | Female | 1 | 41777391 | 41741864 | 99.91% |
|  |  | 2 | 43934472 | 43895938 | 99.91% |
|  |  | 3 | 42383097 | 42341376 | 99.90% |
| Head | Male | 1 | 51631056 | 51544142 | 99.83% |
|  |  | 2 | 46551675 | 46479572 | 99.85% |
|  |  | 3 | 38206399 | 38148536 | 99.85% |
|  | Female | 1 | 54132041 | 54053660 | 99.86% |
|  |  | 2 | 60580208 | 60497298 | 99.86% |
|  |  | 3 | 46450145 | 46377666 | 99.84% |
| Testis | Male | 1 | 45585537 | 45535708 | 99.89% |
|  |  | 2 | 59703504 | 59643374 | 99.90% |
|  |  | 3 | 51146625 | 51079088 | 99.87% |
| Ovary | Female | 1 | 53248762 | 53146800 | 99.81% |
|  |  | 2 | 51869747 | 51785328 | 99.84% |
|  |  | 3 | 57592595 | 57497696 | 99.84% |

**Supplemental Table 5.**
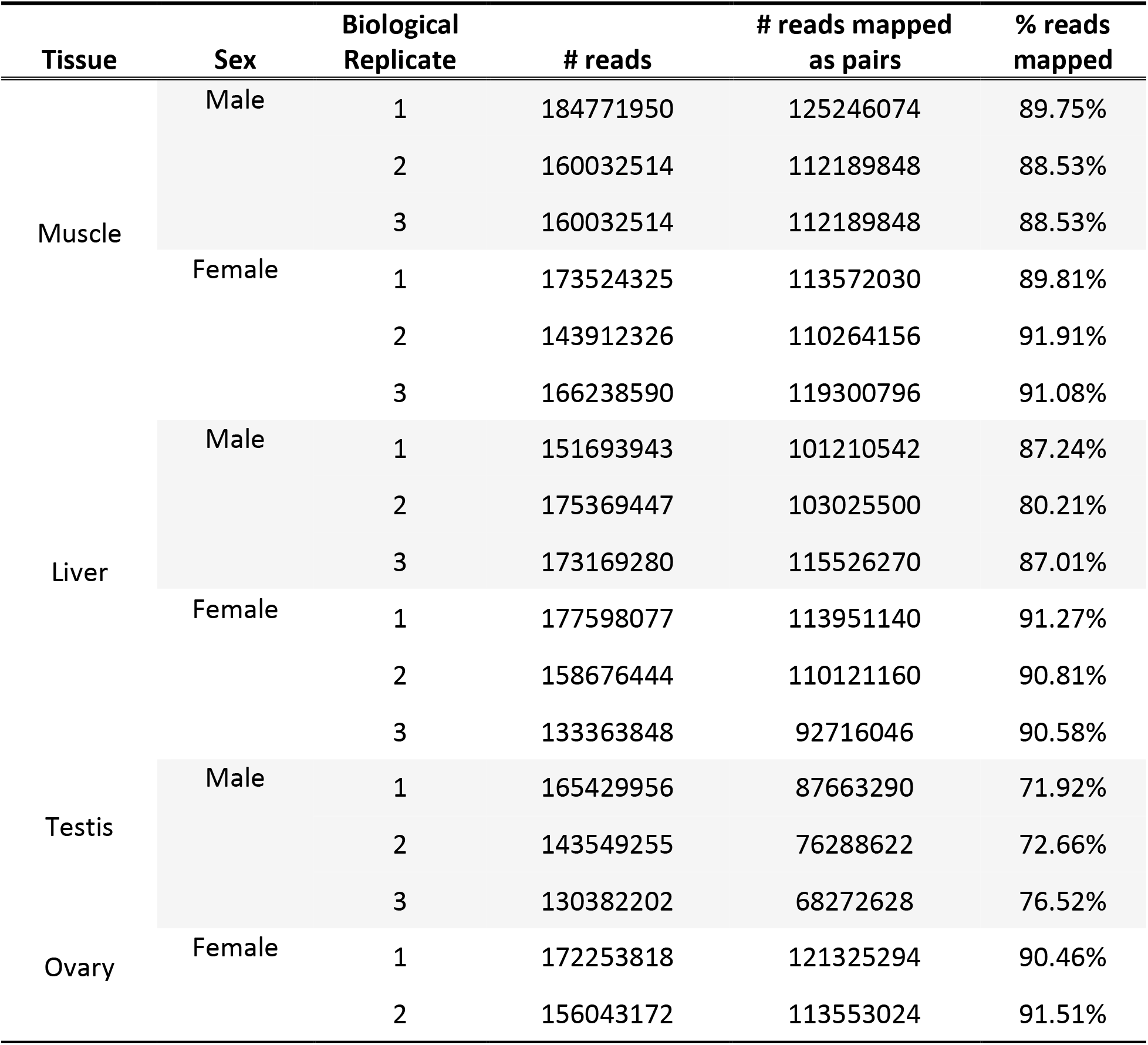
WGBS data for *P. picta*.

| **Tissue** | **Sex** | **Biological Replicate** | **# reads** | **# reads mapped as pairs** | **% reads mapped** |
| --- | --- | --- | --- | --- | --- |
| Muscle | Male | 1 | 184771950 | 125246074 | 89.75% |
|  |  | 2 | 160032514 | 112189848 | 88.53% |
|  |  | 3 | 160032514 | 112189848 | 88.53% |
|  | Female | 1 | 173524325 | 113572030 | 89.81% |
|  |  | 2 | 143912326 | 110264156 | 91.91% |
|  |  | 3 | 166238590 | 119300796 | 91.08% |
| Liver | Male | 1 | 151693943 | 101210542 | 87.24% |
|  |  | 2 | 175369447 | 103025500 | 80.21% |
|  |  | 3 | 173169280 | 115526270 | 87.01% |
|  | Female | 1 | 177598077 | 113951140 | 91.27% |
|  |  | 2 | 158676444 | 110121160 | 90.81% |
|  |  | 3 | 133363848 | 92716046 | 90.58% |
| Testis | Male | 1 | 165429956 | 87663290 | 71.92% |
|  |  | 2 | 143549255 | 76288622 | 72.66% |
|  |  | 3 | 130382202 | 68272628 | 76.52% |
| Ovary | Female | 1 | 172253818 | 121325294 | 90.46% |
|  |  | 2 | 156043172 | 113553024 | 91.51% |

